# Illumina-based Whole *De Novo* Transcriptome Profiling and Diosgenin Biosynthetic pathway of *Tribulus terrestris:* A Medicinal herb

**DOI:** 10.1101/2022.03.21.485112

**Authors:** Parul Tyagi, Rajiv Ranjan

## Abstract

*Tribulus terrestris* L. (Chota gokhuru), an annual herb of the *Zygophyllaceae* family accumulates steroidal-type triterpenoids of diosgenin in root, leaf, fruit, which have been used as a traditional medicine to cure steroidal hormonal diseases such as estrogen, progesterone, and cardiovascular and urogenital. However, to date, the diosgenin biosynthetic pathway of *T. terrestris* is unknown. In this research paper, identification of candidate genes involved in bioactive compounds (diosgenin) was conducted via whole transcriptome profiling of *T. terrestris* using the Illumina *HiSeq* sequencing platform, which generated 482 million reads. RNA-Seq reads were assembled and clustered into 148871 unigenes. NCBI non-redundant (NR) and Swiss-Prot databases search anchored 50% unigenes with the functional annotations based on similarities of sequence. Further, in Gene Ontology and KEGG-KAAS pathways, a total of 126 unigenes were highly expressed in the terpenoids backbone biosynthesis, in which diosgenin pathways (two genes of glycolytic, three genes of mevalonate, two genes of MEP/DOX, one gene of mono-terpenoids, and seven genes of steroidal pathways) were identified. In addition, 9 genes were randomly selected for confirmation through real-time PCR. *T. terrestris* showed the highest homology with the *Citrus sinensis* and *Juglans regia*. 27478 unigenes of transcription factors were assembled, in which top ten transcription factors were highly expressed. A total of 17623 SSR was assembled, amongst them unigenes of di-nucleotide were highly repeated. This research work will improve our understanding of the diosgenin pathway and lay the groundwork for future molecular research on metabolic engineering, as well as raise diosgenin content in *T. terrestris*.

## 1. Introduction

*Zygophyllaceae* is a flowering plant family (trees, shrubs, or herbs) that includes the bean-caper and caltrop. There are around 285 species in 22 genera in the family (Christenhusz et al. 2016). *Tribulus terrestris* (chota gokhuru) is one of the most valuable therapeutic herbs among them. In Asia, it is originated from the sub-tropical regions such as India, China, America, Mexico, and Bulgaria (Akram et al. 2011). It is traditionally used as a medicinal herb for curing numerous diseases, viz. cardiovascular, spermatogenesis, leucorrhoea, sexual dysfunctions, rheumatism, piles, renal, impotency, diuretic, antiseptic, headache, and kidney problems; and also exhibits several biological activities such as cytotoxic, anti-inflammatory, hemolytic, antifungal, and antibacterial properties (Miraj 2016). It is categorized as numerous types of secondary metabolites, namely: saponins, steroids, flavonoids, alkaloids, fatty acids, vitamins, resins, nitrate potassium, aspartic acid, essential oils, and glutamic acid (Vasait and Rajendrabhai 2017). Amongst them, the highest amounts of steroidal saponins were found (Chhatre et al. 2014). In which the “Diosgenin” compound is a steroidal saponin that enhances estrogen and progesterone hormone levels (Qureshi et al. 2014).

At present, the only way to obtain diosgenin is through direct solvent extraction of plant materials, which consumes tones of plant material each year and causes significant environmental pollution. The advent of synthetic biology provides a great opportunity to synthesize diosgenin, but only if a comprehensive list of diosgenin biosynthetic genes is accessible. Despite the therapeutic relevance of diosgenin, its production process remains poorly characterized, particularly in terms of the genes involved in its downstream pathway (Kim et al. 2019). However, there are few molecular research works has done while genomic and transcriptomic data of *T. terrestris* are not available in the NCBI plant database.

Whole genome sequencing of medicinal plants is challenging due to the complexity of their genomes, the cost of sequencing and computational resources. Thus, complete genome sequencing has been performed on notably few medicinal plants. The intricacy of medicinal plants’ genomes, as well as the cost of sequencing and computational resources, makes whole-genome sequencing difficult. Till now, only a few medicinal plants have had their entire genome sequenced. While, transcriptome-based research has been generated by next-generation high-throughput sequencing (Muranaka et al. 2013; Saito 2013). Recent development in sequencing technology has enabled large-scale transcriptome sequencing, allowing for gene expression and functional genomics research.

Whole transcriptome research provides a rapid, high-throughput method for obtaining plant gene function data as well as information on active compounds of biosynthesis and regulation (Wang et al. 2009; Liu et al. 2015). Genome-wide transcription work contributes to our understanding of gene function and the intricate regulatory processes that control gene expression. The majority of genome-wide knowledge is accumulated at the level of gene expression (i.e., variations in mRNA quantity). Although it is commonly considered that each individual gene transcribes identical RNA molecules, one gene can really generate many isoforms via alternative promoters, exons, and terminators. Thus, different RNA molecules (i.e., isoforms) are frequently created during transcription that vary in length and can exhibit considerable differences in function, location, and expression pattern (Kelemen et al. 2013).

The first stage results in the synthesis of 2, 3-oxidosqualene via isoprenoid units derived either from cytosolic mevalonate (MVA) or plastid-targeted 2-C-Methyl-D-erythritol 4-phosphate (MEP). The first-stage genes have been widely studied (Volkman et al. 2005; Vranova et al. 2013). The second stage consists of 10 enzymatic processes that convert 2, 3-oxidosqualene to cholesterol and have been recently described molecularly in plants (Sonawane et al. 2016). However, the genes and enzymes involved in the last stage of the process from cholesterol to diosgenin are mainly unknown, significantly impeding the application of synthetic biology to generate diosgenin in microbes or other biological systems. Multiple oxidations of cholesterol at the C-22, C-16, and C-26 locations are required (Mehrafarin et al. 2010), which may be mediated by particular cytochrome P450s (CYPs).

Comparative transcriptome analysis has aided in the identification of genes involved in the biosynthesis of several key secondary metabolites (He et al. 2012; Kumar et al. 2016). This study gives useful information about *T. terrestris* genes that are up-regulated in response to cholesterol elicitation by differential expression analysis. A set of distinct CYP- and UGT-encoding genes has been proposed for involvement in the downstream stages from cholesterol to diosgenin based on the development of a phylogenetic tree (Supplementary Figure S9) and co-expression analysis with several known upstream genes (Figure 5). Additionally, transcription factors (TFs) involved in diosgenin production were discussed. Thus, our study may serve as a strong foundation for identifying the downstream pathway stages leading to diosgenin, thereby advancing the production of this essential chemical via a synthetic biology strategy. Additionally, a large number of simple sequence repeat (SSR) markers were predicted and developed for diosgenin. These markers can be used in future studies on gene mapping, the development of genetic linkage, the analysis of genetic diversity, and marker-assisted selective breeding of *T. terrestris*.

## 2. Materials and Methods

### 2.1. Collection of plant samples

Root, leaf, and fruit of *Tribulus terrestris* were collected in the month of August from herbal garden of Dayalbagh Educational Institute, Dayalbagh Agra (Tyagi et al. 2021). After cleaning with ultrapure water of all parts and were collected separately, immediately frozen in liquid nitrogen and stored at -80□°C until RNA extraction.

### 2.2. RNA isolation

Total RNA from roots, leaves, and fruits of three replicate were isolated by using *RaFlex Total RNA isolation Kit* (Merck Millipore, Massachusetts, USA) by following the standard protocol defined by the manufacturer. Equimolar (4µl of each sample) concentration was pooled together for RNA-Seq library preparation (Liu et al. 2015). Quality of pooled total RNA were checked by used of 1% formaldehyde denaturing agarose gel and quantity was checked by “Nanodrop 8000 spectrophotometer” (Thermo).

### 2.3. Illumina cDNA library preparation

4µg of total pooled RNA of all parts was taken for paired-end sequencing and prepared cDNA libraries by the use of Illumina *TruSeq Stranded mRNA Sample Preparation Kit* (Illumina, San Diego, California, USA) as per it’s described protocol. mRNA fragments of all samples were enriched in poly-A tailing and assembled into 200bp to 5000bp converted into First-strand cDNA, synthesized (iscript master mix-4µl, iscripts reverse transcriptase-1µl, RNA-4µl, nuclease-free water-11µl) by reverse transcriptase with random hexamer primers. Adopter-ligated libraries (tailing and adapter ligation) were produced for the PCR amplification developed by second-stand generation. DNA High Sensitivity Assay kit on Bioanalyser 2100 (Agilent Technologies, CA, USA) was used for the library quantification and qualification and performed to calculate the mean size of the libraries were 375 base pairs. Nanodrop 8000 spectrophotometer was performed for the quantification of the cDNA libraries (Moraortiz et al. 2016).

### 2.4. De novo assembly and sequencing

RNA-Seq library for cDNA of *T. terrestris* was performed using paired-end (PE) 2×150 bp on Illumina *HiSeq* platform according to house build protocol (Illumina, San Diego, California, USA) (Crawford et al. 2010). The raw data (FASTQ) was decanted as well as removal of adaptor using Trimmomatic v0.30 software (Lindgreen 2012; Manfred et al. 2013; Bolger et al. 2014). Equal or more than N50 values of reads were considered for the calculated per base sequence quality score (QC) or Phred quality score Q≥ 30 was measured (Chen et al. 2018). *De novo* assembly of high-quality reads were accomplished using Trinity v2.1.1 software (Haas et al. 2014) based on De Bruiji graph. Furthermore, the using of CLC Genomic Workbench software (CLC Bio, Boston, MA 02108 USA) for validation of assembled transcript contigs (Wang et al. 2017). Construction of overlapping short reads contigs was assembled with the 30X coverage and depth. Apprize the transcript of GC contains were using a custom-made Perl script (Figure 1).

**Figure 1:**
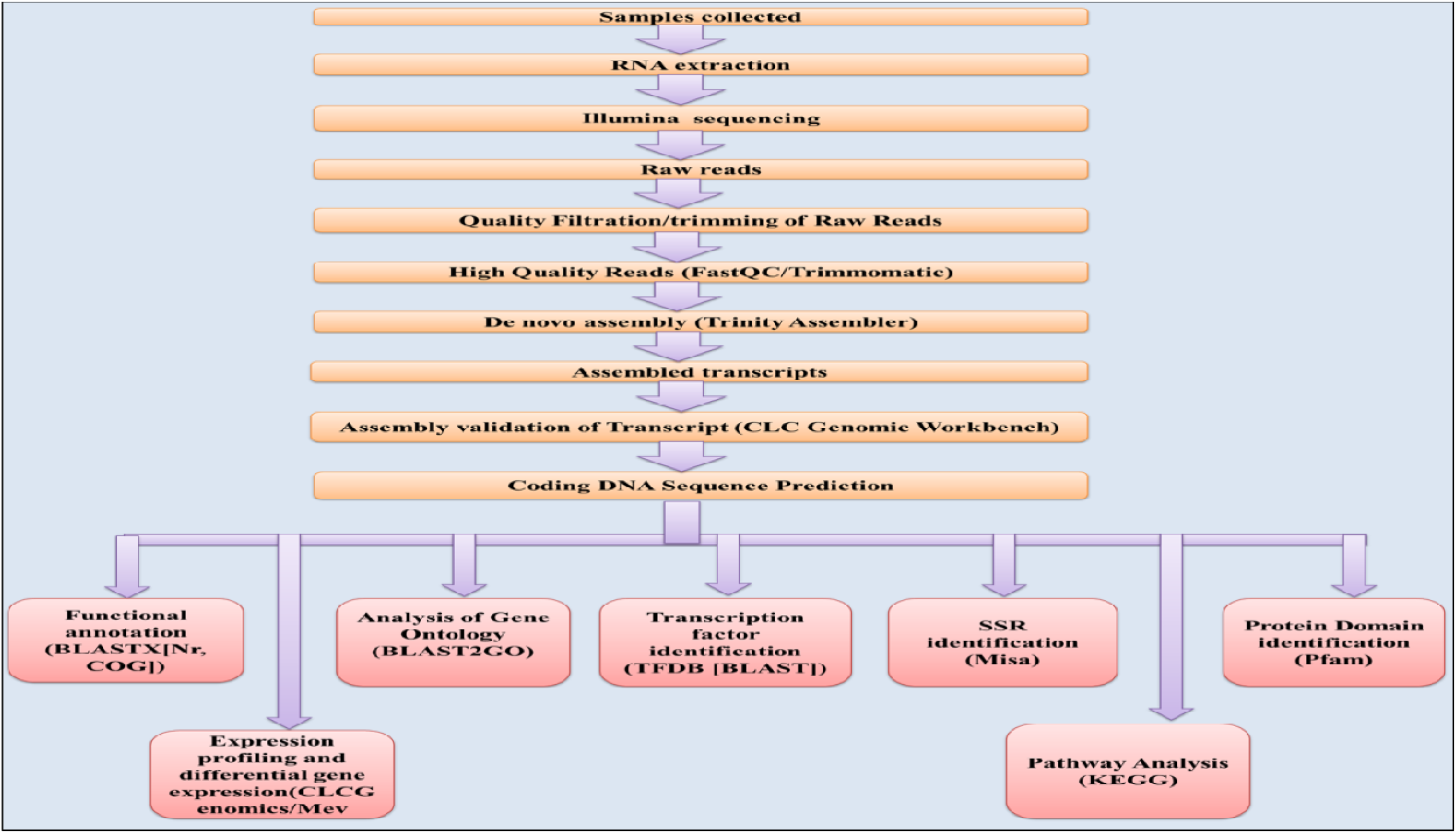
Workflow of Illumina Sequencing platform of all parts of *T. terrestris*.

### 2.5. Functional annotation

Functional annotations of Non-redundant (NR) transcripts are based on several databases; Nr “NCBI non-redundant protein sequences” (https://blast.ncbi.nlm.nih.gov/Blast.cgi) (Altschul et al. 1990), UniProt “The Universal Protein Resource” (http://www.uniprot.org/downloads), Pfam “Protein Families” (http://pfam.xfam.org/), COG “Clusters of Orthologous Groups” (http://www.ncbi.nlm.nih.gov/COG/) databases (Bairoch et al., 2000), GO “Gene Ontology” (http://www.geneontology.org) using *Blast2GO* software (http://www.Blast2go.de/) (Amos et al. 2003; Finn et al. 2014), with an E-value <1e-6. “Kyoto Encyclopedia of Genes and Genomes” (KEGG) (http://www.genome.jp/kegg/) (Moriya et al. 2007; Kanehisa et al. 2014). Transcription factors of assembled unigenes were aligned using BLASTX “Plant Transcription Factor Databases” (http://planttfdb.cbi.pku.edu.cn/). Phylogenetic analysis was performed using the protein sequences of genes involve in diosgenin biosynthesis. The genes were aligned with the same genes of closely related families, and a maximum likelihood phylogenetic tree was built using MEGA X (v.10.2.4) (Trapnell et al. 2010).

### 2.6. Unigenes of Differential expression analysis

The specifically expressed genes were screened from the distinct *T. terrestris* tissues by differential expression analysis. The expression levels of unigenes was calculated using RPKM (Reads per kilobase transcriptome per million mapped reads) for eliminating the influence of gene length and sequencing level on the calculation of gene expression (Moriya et al. 2007), and therefore the calculated results can be directly used to analyze differences in gene expression levels between the different samples by using the statistical comparison method (Audic et al. 1997). The obtained results of multiple hypothesis testing were corrected by False Discovery Rate (FDR) control method, and the ratio of RPKMs was converted to the fold-change in the expression of each gene in two samples simultaneously. Afterwards, the significance differentially expressed genes were screened by the threshold of FDR≤0.001 and the absolute value of log2Ratio≥1, and then were mapped to database for pathway enrichment analysis.

### 2.7. Analysis of transcription factors (TFs)

To assess the transcription factor (TFs) families represented in the *T. terrestris* transcriptome, the open reading frame (ORF) of each unigenes was determined using the software getorf (EMBOSS:6.5.7.0) (Mistry et al. 2013). Using BLASTX (e-value 1e-6), we aligned these ORFs to all the TF protein domains in the plant transcription factor database (*PlnTFDB*) (Unamba et al. 2015).

### 2.8. Analysis of Quantitative Real-Time (qRT-PCR)

To validate the RNA-Seq results, qRT-PCR analysis was performed using 96 real-time PCR systems (Bio-Rad, USA) equipped with a SYBR^®^ Premix Ex Taq™ kit (Agilent Aria (v1.5) Software) (Takara, China). IDT (https://eu.idtdna.com/pages) software was used to design for QRT-PCR gene primers (SupplementaryTable S4). Successive qRT-PCR experiments were done using three biological and three technical replicates, and melting curve analysis was performed following every amplification to confirm product specificity. To normalize, the alpha-tubulin gene from *T. terrestris* was used as an internal reference gene, and each sample was divided into three technical duplicates. The relative expression levels of each gene were calculated using the 2^−ΔΔCt^ method (Livak et al. 2001).

### 2.9. *SSRs* identification

Micro Satellite software Perl script “MISA” (http://pgrc.ipk-gatersleben.de/misa/) was used to identify Simple Sequence Repeats (SSRs), in which unigenes were used as reference data and transcript contigs were searched for SSRs. The sequence was originally prepared and mined for SSR motifs sequences on both ends (dimer to hexamer nucleotide) with a length of 150 bp and these were retained for di-, tri-, tetra-, and hexa-nucleotide repeats are mentioned in Supplementry Table S5 (Moraortiz et al. 2016).

## 3. Results

### 3.1. De novo assembly of RNA sequencing

Genomic research in non-model plants can now be facilitated by NGS technologies such as novel gene discovery and tissue-specific expression analyses as well as the development of sequence-based molecular marker resources (Grabherr et al. 2013). To define the transcriptome profile of *T. terrestris*, we sequenced nine cDNA libraries generated from root, leaf, and fruit tissues in three biological replicates using the Illumina *Hiseq*2000 platform’s paired-end (PE) sequencing. After deleting adaptors, poly-A tails and primer sequences, as well as short (150 bp) and low quality sequences, a total of ∼7.9 GB of clean data were retrieved. A total of 410 million raw readings were generated, as well as pooled tissues. There were 14150090 (29.33 %) duplicate reads obtained, and 112182 transcript contigs were constructed from all clean reads. Supplementary Figure S1 and Table 1 demonstrate that 28.62 % (34975930), 28.49 % (34820562), 21.68 % (26488197), and 21.90 % (25918968) of ATGC composition percentages were obtained from all areas of *T. terrestris*. To eliminate duplicated sequences, the high-quality reads and all isoforms were assembled using the Trinity programme (Perteal et al. 2003) and the TGI clustering tool (TGICL) (Lulin et al. 2012).

**Table 1:** Transcriptome data of whole herbs parts (root, fruit, and leaf) of *Tribulus terrestris*.

| **Samples of *T. terrestris*** | **Raw data** |
| --- | --- |
| Total reads | 48239004 |
| Total duplicate reads | 14150090 (29.33%) |
| Total bases | 7032788672 |
| Total numbers of transcripts | 245069 |
| Total transcriptomics length in bps | 179930901 |
| Average transcript length in bps | 734.20 |
| Maximum length of transcripts (>5000) bps | 462 |
| Minimum length of transcripts (200-300) bps | 88552 |
| N50 | 1254 |
| Total numbers of unigenes | 200471 |
| Total size of all unigenes in bps | 122203657 |
| Average unigenes length in bps | 609.58 |
| Maximum length of unigenes bps | 12380 |
| Minimum length of unigenes bps | 201 |
| N50 | 956 |
| GC content | 52407165 (42.89%) |
| Depth of unigenes | 122203657 (58%) |
| Total number of CDS | 112182 |
| Total size of all CDS in bps | 69264741 |
| Average CDS length in bp | 617.43 |
| Maximum CDS length in bps | 11637 |
| Minimum CDS length in bps | 297 |
| Total annotated CDS hits with BLASTX | 100753 |
| Total without annotated CDS hits with BLASTX | 11429 |
| N50 | 684 |
| Unigenes in Uniprot databases | 58746 |
| Unigenes in COG database | 37292 |
| Unigenes in GO database | 96899 |
| Unigenes in Pfam database | 35989 |
| Unigenes with KEGG database | 7721 |

Supplementary Figure S2 shows that a total of 12380 unigenes with a minimum length of 200-300 bp (maximum unigenes) and 201 unigenes with a maximum length of 900-1000 bp (minimum unigenes) were generated. G+C percentage of [40251760 (43.34%)] and depth contents of [122203657 (65%)] unigenes were obtained are shown in Supplementary Figure S3. A total of 200471 unigenes were identified, with an average N50 length of 956 bp (nucleotide) and an average length of 609.58 bp. Among them, a total of 691 up-regulated and 100 down-regulated unigenes were annotated as being involved in diosgenin metabolic biosynthetic pathways. All transcriptome data from whole de novo sequencing are shown in (Figure 1) and Table 1, which were prepared using a custom-written Perl script (Gene Ontology 2004; Nakasugi et al. 2014; Lehnert et al. 2014). Correlation indices between repeated samples were >0.9, demonstrating the validity of the Illumina sequencing data. Raw reads of *T. terrestris* was generated using Illumina *Hiseq*2000 sequencing and submitted in the National Center for Biotechnology Information (NCBI) Sequence Read Achieve (SRA) database under the accession number PRJNA786412.

### 3.2. Functional annotation

The functional characterization of transcriptome data gives a comprehensive view of biological processes, molecular functions and cellular components, and the abundance of biosynthetic pathways. *T. terrestris* (as a non-model plant), transcripts were aligned with six public protein databases to provide the most accurate annotations. We searched all assembled unigenes against the Non-redundant (Nr); using the BLASTx programme with an E-value of 1e-6. Uniprot is a comprehensive database of protein sequences and annotations; “Kyoto Encyclopedia of Genes and Genomes (KEGG)” is a database resource for deriving high-level functions and utilities of biological systems, such as the cell, the organism, and the ecosystem, from molecular-level information, particularly large-scale molecular datasets generated by genome sequencing; The purpose of Pfam is to provide an exhaustive and accurate classification of protein families and domains; Gene Ontology (GO) is a statement about the function of a particular gene; and Clusters of Orthologous Groups (COG) is an attempt at phylogenetic classification of the proteins encoded in 21 complete genomes of bacteria, archaea, and eukaryotes. A total of 200471 unigenes were functionally assembled, of which 37292, 42256, 35989, and 6940 unigenes from COG, uniprot, pfam, and KEGG databases, respectively, shown Supplementary Figure S4. Amongst them, 5185 unigenes are obtained in common which are found in all databases. The results of BLASTX searches against several databases identified clusters of orthologous proteins (COGs) and their annotations are shown in (Figure 2) and Supplementary File S1, and categorized genes based on their orthologous homology. Based on the Nr database, the E-value distributions indicated that 1000753 of the matched unigenes ranged from 1e-6 to 1e-100 and show a maximum number of unigenes in *Quercus suber, Theobroma cacao, Vitis vinifera, Juglans regia*, and *Citrus sinensis*. Amongst them, *T. terrestris* showed higher similarity with the *Quercus suber* plant. A total of 200471 unigenes were displayed in the similarity distribution are shown (Supplementary Figure S5).

**Figure 2:**
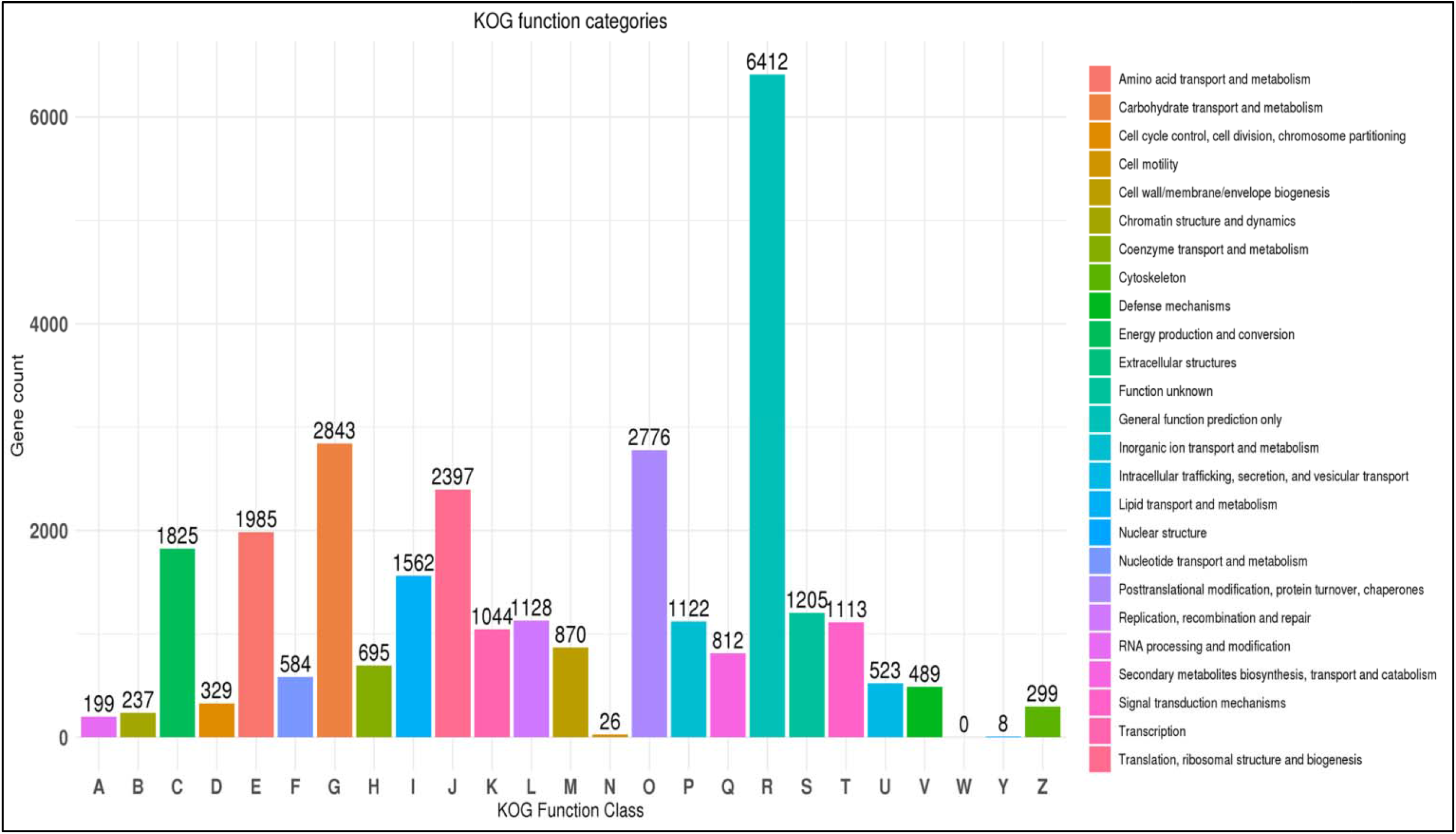
Function annotation classifications: Cluster Orthologous Groups (COG) classification of transcripts into 25 categories; database of *T. terrestris*.

Gene ontology (GO) has been widely employed for functional analysis and inferring the biological significance of genomic and transcriptome data (Fujita et al. 2006). Three major domains were identified in the GO analysis including biological processes (BP), cellular components (CC), and molecular functions (MF). When the Gene Ontology (GO) classification system was used to categorized gene functions, 46669 unigenes were classified into 60 functional categories, which are listed in Supplementry File S2. In which, 33258 of BP domain (operations or sets of molecular events with a defined start and end that are required for the proper functioning of integrated living units such as cells, tissues, organs, and organisms); 26597 of CC domain (components of a cell or its extracellular environment); and 37044 of MF domain (the elemental activities of a gene product at the molecular level, such as binding or catalysis) are shown in (Figure 3) and Supplementary Table S1. We matched the annotated unigenes to the reference biological pathways in the KEGG database to better understand the roles of distinct metabolic pathways in *T. terrestris*. Supplementary Figure S6 shows a total of 35989 pfam domains are identified in whch highest number of Pkinase and Pkinase_Tyr enzymes are obtained. A total of 112182 CDS that were assembled and analyzed against the KEGG databases using BLASTX with the threshold bit-score value of 60 (default). In *T. terrestris*, a total of 151 pathways were obtained, including 11 metabolic routes, 4 genetic information processing pathways, 3 environmental information processing pathways, 5 cellular process pathways, 10 organismal system pathways, 12 human illness, and 3 brite hierarchy pathways, etc.

**Figure 3:**
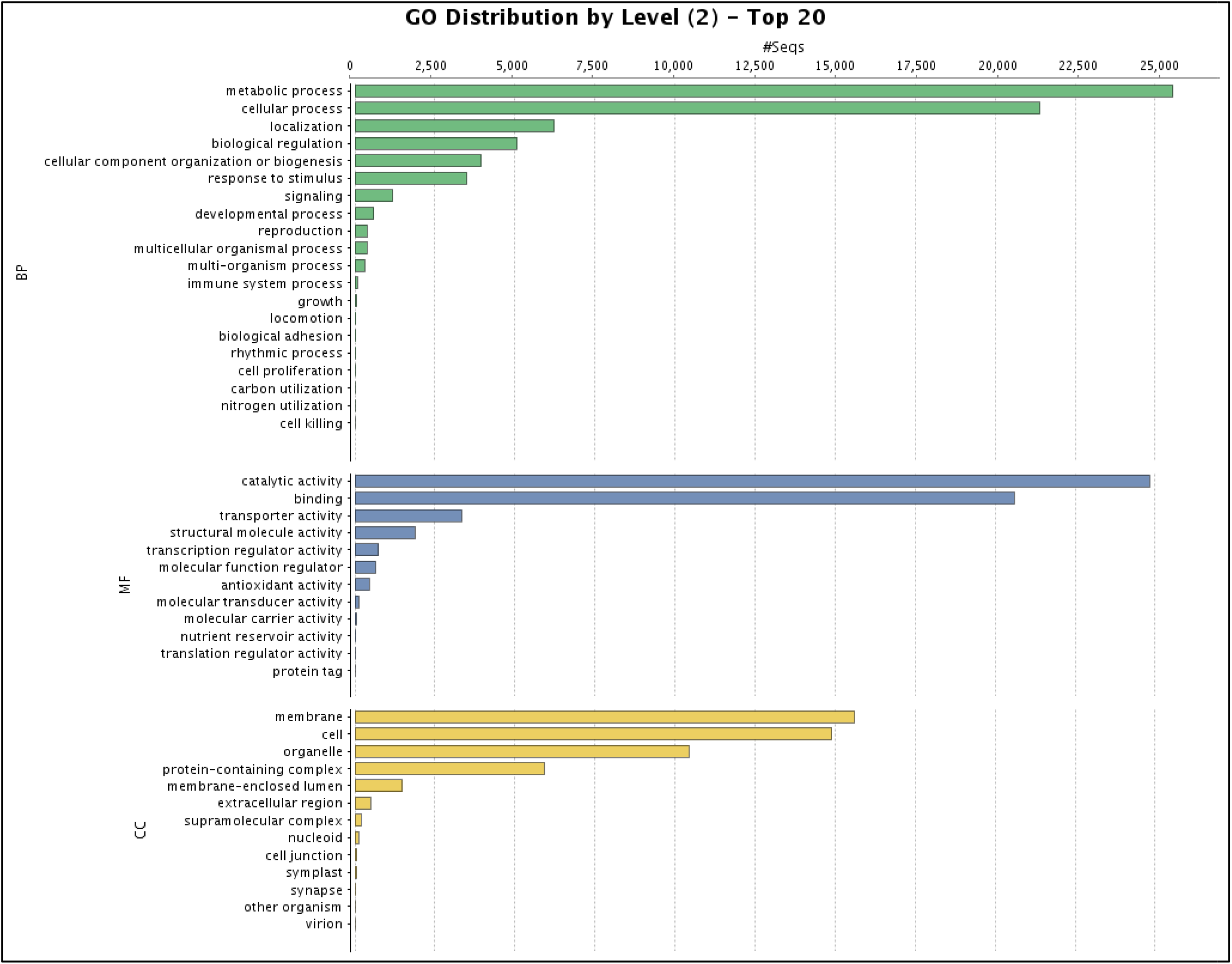
Function annotation classifications: Gene Ontology (GO) classification summarized into, cellular component, biological process and molecular function categories of *T. terrestris*.

KEGG annotations provide context for continuing metabolic processes within an organism, providing for a better understanding of the biological function of the transcripts (Kanehisa et al. 2014). The distribution of 17,786 unigenes into six major categories and 45 subcategories is depicted in (Figure 4) and Supplementary Table S2. The enzymes with assigned roles in 3567 CDS metabolic pathways in KEGG are illustrates Supplementary Figure S7. Among these unigenes, 126 CDS encode important enzymes involved in the synthesis of terpenoid backbones (31 unigenes), monoterpenoids (3 unigenes), diterpenoids (21 unigenes), sesquiterpenoids and triterpenoids (7 and 7 unigenes), and various terpenoid-quinone complexes (50 unigenes). There are five unigenes involved in tropane, piperidine, and pyridine biosynthesis and six unigenes involved in isoquinoline alkaloid biosynthesis. Exactly 102 unigenes were related with the flavonoid biosynthesis pathway, which included the phenylpropanoid (49 unigenes), flavonoid (49 unigenes), flavone, and flavonol production pathways (2 unigenes), etc.

**Figure 4:**
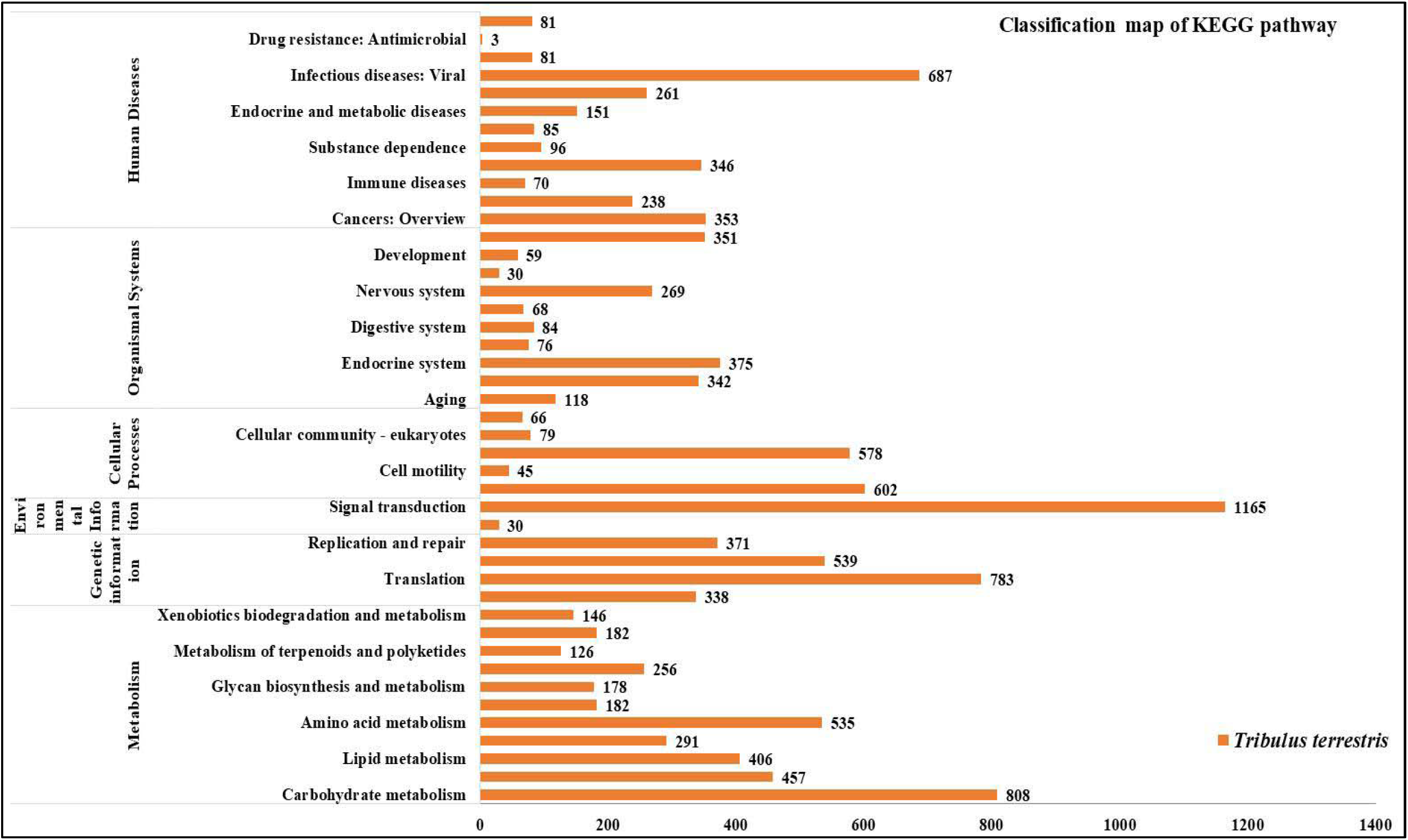
Function annotation classifications: Kyoto Encyclopedia of Genes and Genomes (KEGG) classification details of six main categories, I: Metabolism, II: Genetic information processing, III: Environmental information processing, IV: Cellular processes, V: Organismal systems, (VI) Human diseases of root, fruit, and leaf tissues of *T. terrestris*.

**Figure 5:**
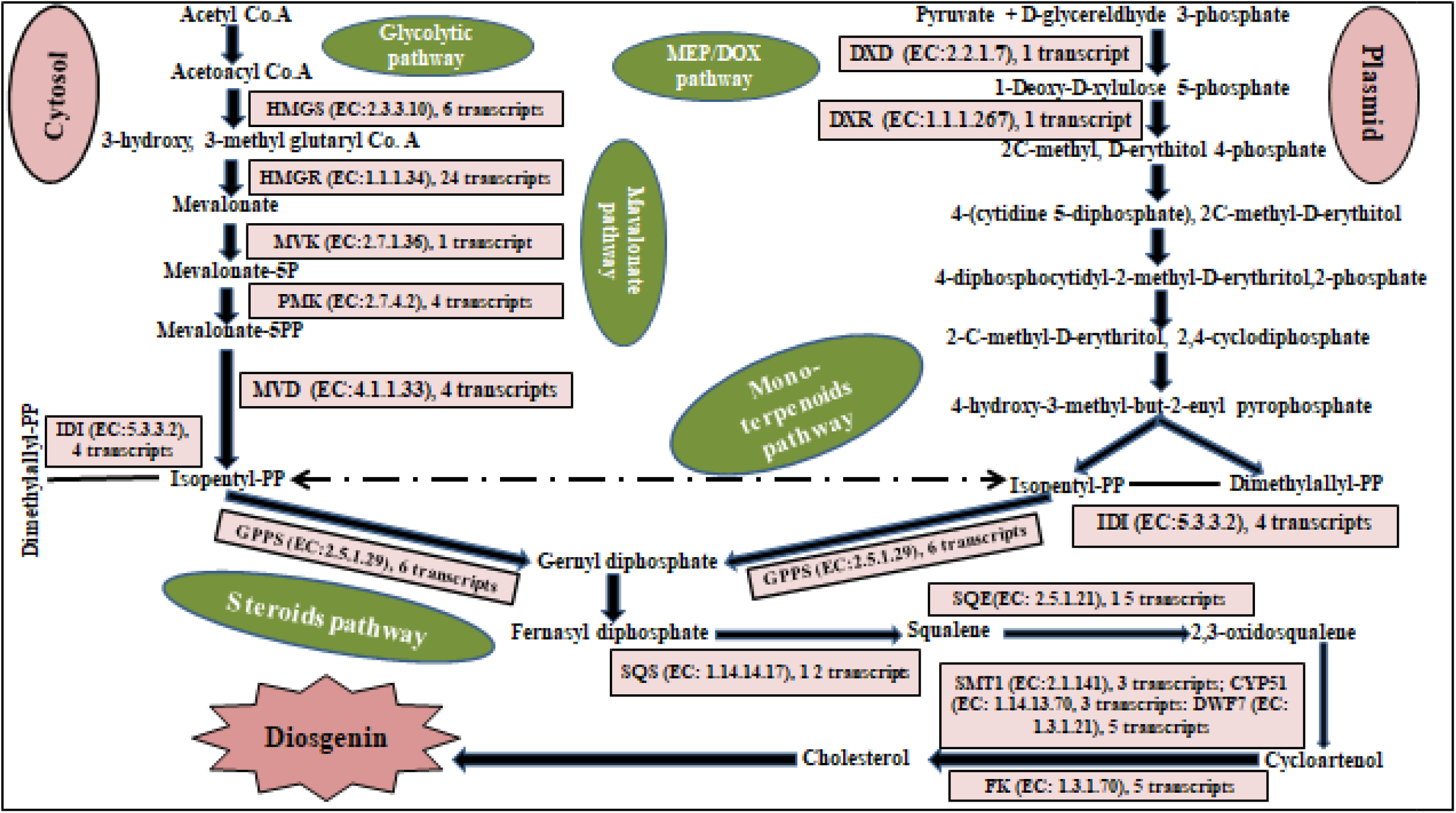
Schematic representation of the potential key genes of diosgenin biosynthesis pathway of diosgenin. *Alpha Actin*: Housekeeping genes; *HMGS*: hydroxymethyl-glutaryl-CoA synthase; *HMGR*: hydroxyl methylglutarylCoA reductase; *MVK*: mevalonate kinase; *PMK*: Phosphomevalonate kinase; *MVD*: Mevalonate diphosphate decarboxylase; *DXD*: 1-deoxy-D-xylulose-5-phosphate synthase; *DXR*: 1-deoxy-D-xylulose-5-phosphate reductoisomerase; *IDI*: Isopentenyl-diphosphate delta-isomerase; *GPPS*: Geanylgeranyl diphosphate synthase; *SQE*: Squalene epoxidase; *SQS*: Squalene synthase; *DWF7*: 7-dehydrocholesterol reductase; *SMT1*: Sterol 24-C methyltransferase; *FK*: Delta 14-sterol reductase.

In order to have better information of diosgenin biosynthesis, we identified 2116 unigenes involved in **Metabolism (KO09100):** carbohydrate metabolism (ko09010, 808), amino acid metabolism (ko09105, 535), nucleotide metabolism (ko09104, 291), energy metabolism (ko09102, 457), lipid metabolism (ko09103, 406), metabolism of other amino acids (ko09106, 182), glycan biosynthesis and metabolism (ko09107, 178), metabolism of cofactors and vitamins (ko09108, 256), metabolism of terpenoids and polyketides (ko09109, 126), biosynthesis of other secondary metabolites (ko09110, 182), xenobiotics biodegradation and metabolism (ko09111, 146); **Genetic Information Processing (KO09120):** transcription (ko09121, 338), translation (ko09122, 783), folding, sorting and degradation (ko09123, 539), replication and repair (ko09124, 371); **Environmental information processing (KO09130):** membrane transport (ko09131, 30), signal transduction (ko09132, 1165), signalling molecules and interaction (ko09133, 0); **Cellular processes (KO09140):** transport and catabolism (ko09141, 602), cell motility (ko09142, 45), cell growth and death (ko09143, 578), cellular community-eukaryotes (ko09144, 79), cellular community-prokaryotes (ko09145, 66); **Organismal systems (KO09150):** aging (ko09149, 118), immune system (ko09151, 342), endocrine system (ko09152, 375), circulatory system (ko09153, 76), digestive system (ko09154, 84), excretory system (ko09155, 68), nervous system (ko09156, 269), sensory system (ko09157, 30), development (ko09158, 59), environmental adaptation (ko09159, 351); **Human diseases (KO09160):** cancers: overview (ko09161, 353), cancers: specific types (ko09162, 238), immune diseases (ko09163, 70), neurodegenerative diseases (ko09164, 346), substance dependence (ko09165, 96), cardiovascular diseases (ko09166, 85), endocrine and metabolic diseases (ko09167, 151), infectious diseases: bacterial (ko09171, 261), infectious diseases: viral (ko09172, 687), infectious diseases: parasitic (ko09174, 81), drug resistance: antimicrobial (ko09175, 3), drug resistance: antineoplastic (ko09176, 81); **Brite hierarchies (KO09180):** protein families: metabolism (ko09181, 1243), protein families: genetic information processing (ko09182, 5122), protein families: signaling and cellular processes (ko09183, 1126), etc. pathways based on the KEGG database. A total of 11,145 unigenes encoding key enzymes involved in diosgenin biosynthesis including transferases (4011), oxidoreductases (2938), hydrolases (2432), ligases (763), isomerases (593), lyases (408), etc these were screened out in Supplementary Figure S7. Using the “Fragments Per Kilobase of transcript per Million FPKM” method, these data allowed to revealed that the genes encoding enzymes which involved in diosgenin production. To further understand their roles in the production of active compounds in *T. terrestris*, unigenes participating in these pathways should be found further (Figure 5).

### 3.3. Differentially expressed gene (DEG) analysis

The edgeR tool was used to analyse tissue-specific gene expression in order to evaluate a better understanding of the key potential regulators involved in steroidal saponin synthesis. The clean reads were mapped back onto the assembled unigenes by using Trinity software to assess DEGs between various tissues of *T. terrestris*. The value of fragments per kilobase million (FPKM) was determined for each unigene of *T. terrestris* tissue. DEGs were identified using a q value ≤□0.005 and log2 (fold change) ≥1 (Storey, et al., 2010). Comparing transcripts in different tissues involved in the diosgenin biosynthesis pathway pairwise led to the identification of 783 DEGs in the total root, fruit, and leaf of *T. terrestris* (687 up-regulated and 97 down-regulated). A total of 105 isoform were expressed in all parts of *T. terrestris* included ***HMGS* (5 isoform:** unigene_30737_cds_14246, unigene_33731_cds_15931, unigene_33731_cds_15932, unigene_33731_cds_15933, unigene_45825_cds_24009); ***HMGR* (25 isoform:** unigene_10860_cds_4481, unigene_10860_cds_4482, unigene_10870_cds_4484, unigene_10878_cds_4485, unigene_10879_cds_4486, unigene_35461_cds_17023, unigene_35461_cds_17024, unigene_43929_cds_22676, unigene_43929_cds_22677, unigene_44874_cds_23351, unigene_82481_cds_57837, unigene_82482_cds_57838, unigene_82484_cds_57841, unigene_82485_cds_57843, unigene_82486_cds_57847, unigene_82487_cds_57851, unigene_82488_cds_57856, unigene_90490_cds_66302, unigene_90492_cds_66303, unigene_90492_cds_66304, unigene_90492_cds_66305, unigene_90493_cds_66306, unigene_90493_cds_66307, unigene_90493_cds_66308, unigene_131048_cds_98880); ***MVK* (1 isoform:** unigene_82578_cds_57949); ***PMK* (4 isoform:** unigene_5926_cds_2619, unigene_9829_cds_4162, unigene_38555_cds_18989, unigene_38555_cds_18990); **MVD (4 isoform:** unigene_42875_cds_21937, unigene_53200_cds_29985, unigene_59157_cds_35079, unigene_77772_cds_52943); ***IDI* (4 isoform:** unigene_15193_cds_6303, unigene_32889_cds_15450, unigene_59108_cds_35030, unigene_116224_cds_93504); ***DXD* (9 isoform:** unigene_27514_cds_12458, unigene_42741_cds_21868, unigene_42742_cds_21869, unigene_60891_cds_36625, unigene_65190_cds_40617, unigene_65190_cds_40618, unigene_65191_cds_40619, unigene_65191_cds_40620, unigene_70011_cds_45258); ***DXD* (3 isoform:** unigene_90670_cds_66485, unigene_90670_cds_66486, unigene_90672_cds_66487); ***GPPS* (6 isoform:** unigene_3738_cds_1605, unigene_3738_cds_1606, unigene_68239_cds_43581, unigene_84044_cds_59611, unigene_105751_cds_84117, unigene_173740_cds_108466); ***SQS* (12 isoform:** unigene_28533_cds_12992, unigene_28533_cds_12993, unigene_83903_cds_59474, unigene_83903_cds_59475, unigene_83904_cds_59476, unigene_83904_cds_59477, unigene_92926_cds_69186, unigene_92926_cds_69187, unigene_126178_cds_97559, unigene_139109_cds_101535, unigene_141551_cds_101999, unigene_159016_cds_106054); ***SQE* (15 isoform:** unigene_15216_cds_6322, unigene_15216_cds_6323, unigene_15216_cds_6324, unigene_15223_cds_6327, unigene_58318_cds_34332, unigene_58318_cds_34333, unigene_58319_cds_34334, unigene_58320_cds_34335, unigene_58320_cds_34336, unigene_89105_cds_64945, unigene_89105_cds_64946, unigene_96074_cds_72753, unigene_96076_cds_72754, unigene_96076_cds_72755, unigene_154475_cds_104744); ***SMT1* (3 isoform:** unigene_45046_cds_23473, unigene_45047_cds_23474, unigene_45048_cds_23475); ***CYP51* (3 isoform:** unigene_18413_cds_7981, unigene_33070_cds_15566, unigene_86484_cds_62033); ***FK* (5 isoform:** unigene_12495_cds_5085, unigene_12495_cds_5086, unigene_50584_cds_27729, unigene_50584_cds_27730, unigene_66509_cds_41877); ***DWF7* (6 isoform:** unigene_12809_cds_5207, unigene_12809_cds_5208, unigene_58874_cds_34852, unigene_58875_cds_34853, unigene_58877_cds_34857, unigene_140593_cds_101815) were obtained in diosgenin biosynthetic pathway analysis.

Meanwhile, the top 20 significant level of gene ontology in *T. terrestris* were analyzed based on the FDR □ ≤ □ 0.05 including **“biological process”:** metabolic process, cellular process, localization, biological regulation, response to stimulus, cellular component organization or biogenesis, signaling, multicellular organismal process, developmental process, multi-organism process, reproduction, immune system process, locomotion, growth, biological adhesion, cell proliferation, behavior, rhythmic process, pigmentation, cell killing; **“molecular function”:** catalytic activity, binding, structural molecular activity, transporter activity, transcription regulator activity, molecular function regulation, antioxidant activity, molecular carrier activity, nutrient reservoir activity, cargo receptor activity, toxin activity, transcription regulator activity; **“cellular process”:** cell, membrane, organelle, protein-containing complex, membrane-enclosed lumen, extracellular region, super molecular complex, virion, synapse, other organism, cell junction, nucleoid, and symplast were found in significant enrichment are shown in (Figure 3) and Supplementary Table S1 In addition to common primary metabolic pathways, enriched secondary metabolic pathways, such as terpenoid and diosgenin biosynthesis, were identified in *T. terrestris* tissues, indicating the herb’s possible number of different of secondary metabolites.

### 3.4. Identification of Transcription Factors

Transcription factors (TFs) attach to certain cis-regulatory components of promoter regions and play an important role in gene expression, plant secondary metabolism, and plant response to environmental stress. In numerous plant species, TF families such as ARF, bHLH, bZIP, MYB, NAC, and WRKY have been implicated in secondary metabolite regulation, as well as abiotic and biotic stress responses (Patra et al. 2013; Yin et al. 2017). A total of 21026 unigenes of *T. terrestris* were annotated as transcription factors (TFs), and they were grouped into 10 transcription factor families including “basic helix-loop-helix (bHLH)” (2779), “NAC (NAM, ATAF and CUC)” (1990), MYB related family (highly conserved) (1984), “APETALA2/Ethylene Responsive Factor (AP2/ERF)” (1578), “Zinc finger C2H2” (1398), WRKY (1217), “Cysteine3Histidine C3H” (1967), “Myeloblastosis MYB” (1010), “Basic leucine zipper Bzip” (1011), B3 (973), etc were highly expressed. Members of the WRKY, bHLH, and AP2/ERF families have been found to play a role in regulating the biosynthesis of terpenoids, alkaloids, and steroids are shown in Supplementary Figure S8 (Paul et al. 2017; Schluttenhofer et al. 2015). Amongst them, the bHLH, NAC, and MYB transcription family were founded in most abundant number. All of these TFs are contributing to the gene expression profiling of T. terrestris by binding to cis-regulatory specific regions in the promoters of their target genes, allowing for further research into their regulatory actions in diosgenin production.

### 3.5. qRT-PCR validation of selected genes

To validate the Illumina sequencing expression profiles, we performed qRT-PCR on nine chosen genes associated to triterpene diosgenin biosynthesis, as shown in (Figure 6). Consistent with the Illumina data, most of these genes were widely expressed in *T. terrestris* root, fruit, and leaf tissues, with acetyl-CoA acetyltransferase, hydroxymethylglutaryl-CoA synthase, hydroxymethylglutaryl-CoA reductase, and squalene epoxidase (SQE) genes being the most abundant. The fold variations in expression were likewise similar to the RNA-seq data. The qRT-PCR results show that the RNA-seq data from this analysis were reliable. A total of 14 genes were found in transcriptome data analysis of root, fruit, and leaf of *T. terrestris* which are related to diosgenin biosynthesis in that 9 genes were selected for validation by Quantitative Real-Time Polymerase Chain Reaction (qRT-PCR). Supplementary Table S4 lists the primers for these selected genes that were used in the qRT-PCR analysis.

**Figure 6:**
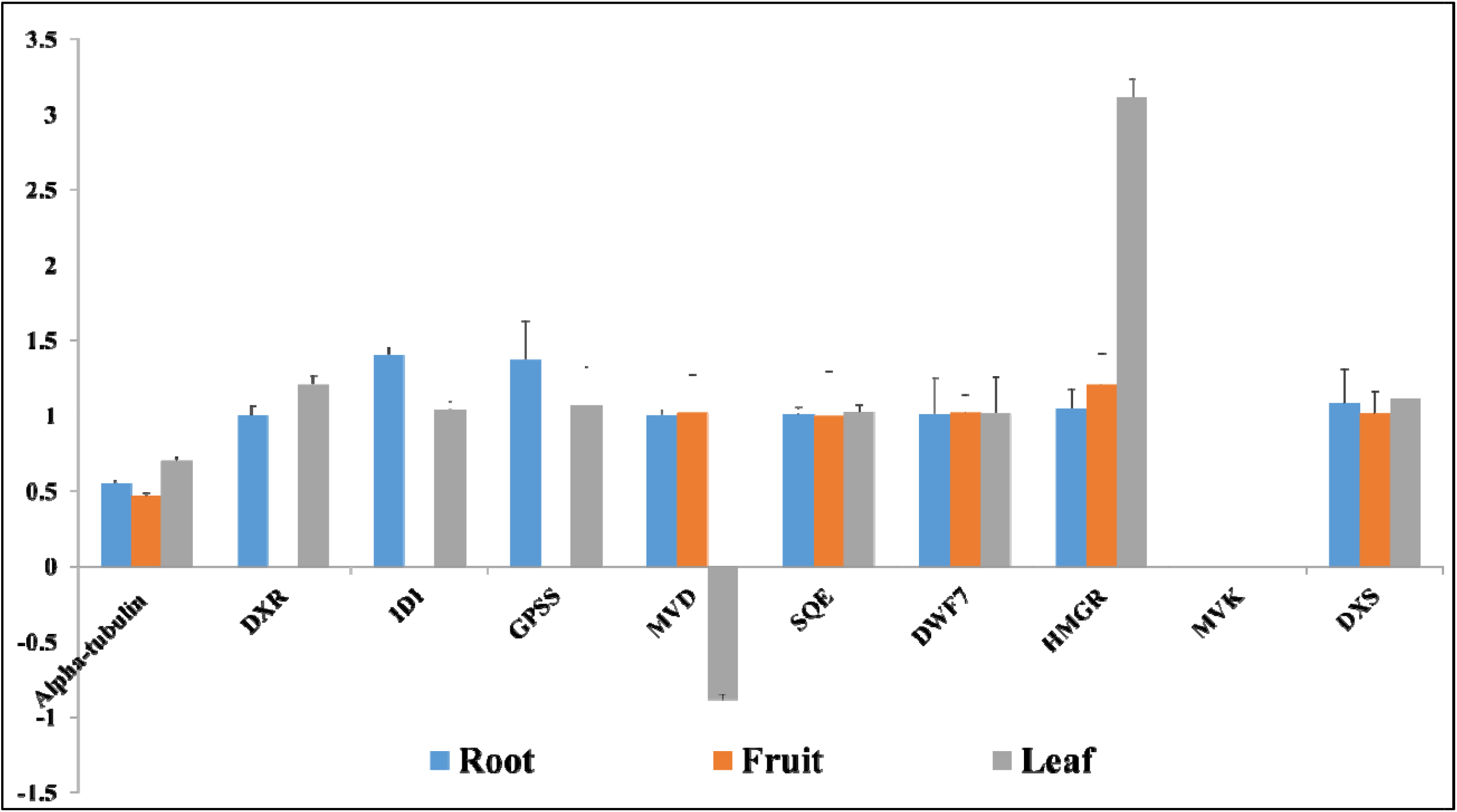
qRT-PCR validation of selected nine genes related to diosgenin biosynthesis pathway. X-axis represents genes and Y-axis is the relative fold change in gene expression by considering root, fruit, and leaf tissues of *T. terrestris*.

The study of metabolic pathways aims to learn about the relationships between genes in a given pathway and their biological activities. Tri-terpenes are synthetized from a five-carbon isoprene unit through the cytosolic MVA pathway. Steroids diosgenin are composed of six isoprene units and are derived from the C-30 hydrocarbon precursor of squalene. It is synthesized from isopentenyl diphosphate *(IPP)* via the MVA pathway. All genes encoding the enzymes associated with the up-regulate and down-regulate regions of steroids diosgenin biosynthesis were successfully detected in the *T. terrestris* transcriptome. Their expression value was monitored in three biological triplicates along with their mean values. Hydroxymethylglutaryl-CoA reductase (*HMGR)* gene in leaf, geanylgeranyl diphosphate synthase (*GPPS)* and Isopentenyl-diphosphate delta-isomerase *(IDI)* genes in root are highly expressed. While, 1-deoxy-D-xylulose-5-phosphate reductoisomerase *(DXR), IDI*, and *GPPS* genes absent in fruit. Mevalonate kinase (*MVK)* gene are absent in root, leaf, and fruit are shown in (Figure 6), Supplementary Table S4. Most of unigenes are related to mevalonate and steroidal pathways were specifically up-regulated in the leaf and root. Supplementary File S2, list of transcripts related to triterpenoid diosgenin backbone biosynthesis are shown in (Figure 5).

### 3.6. Identification of SSRs

Simple sequences repeat (SSR) and microsatellites are the most important for the molecular marker, which is used for gene mapping, molecular breeding, and genetic diversity. These are the tandem repeats of nucleotide motifs of the sizes 2-6 bp and they are highly polymorphic and are ubiquitously present in all the known genomes. Thus SSR were identified from assembled transcript sequences with the MISA perl. SSRs having a flanking of 150 bp (upstream as well as downstream) were fetched with in-house python script which can be used for primer designing. A total number of 15551 SSRs loci with di-, tri-, tetra-, penta- and hexanucleotide repeats were recognized in which, 9292 of di-nucleotide, 6580 of tri-nucleotide, 146 of tetra-nucleotide, 4 of penta-nucleotide motifs were found and the most abundant of repeats including AC/GT (770), AG/CT (7842), AT/AT (1492), CG/CG (21), AAC/GTT (380), AAG/CTT (3065), AAT/ATT (335), ACC/GGT (407), ACG/CGT (175), ACT/AGT (90), AGC/CTG (694), AGG/CCT (341), ATC/ATG (1203), CCG/CGG (445), AAAC/GTTT (24), AAAG/CTTT (105), AAAT/ATTT (85), AACC/GGTT (5), AAGC/CTTG (6), AAGG/CCTT (6), AAGT/ACTT (1), AATC/ATTG (6), AATG/ATTC (6), AATT/AATT (18), ACAG/CTGT (5), ACAT/ATGT (20), ACCG/CGGT (1), ACGG/CCGT (1), ACTC/AGTG (11), ACTG/AGTC (1), AGAT/ATCT (22), AGCC/CTGG (1), AGCG/CGCT (2), AGGC/CCTG (1), ATCC/ATGG (1), ATGC/ATGC (16), AAAAT/ATTTT (1), AAACG/CGTTT (1), AAATC/ATTTG (4), AACGG/CCGTT (1), AATGT/ACATT (1), ACACC/GGTGT (6), ACAGC/CTGTG (1), ACCAG/CTGGT (1), ACGAT/ATCGT (1), ACTGC/AGTGC (1) and AGCTC/AGCTG (1), etc respective motif repeats were obtained.

## 4. Discussion

Many technologies have been used to analyze and quantify the transcriptome of model or non-model organisms, such as Arabidopsis, rice, radish (*Raphanus sativus* L.), and *Haloxylon ammodendron*; as such, these techniques are vital to elucidate the complexity of growth and development of organisms. For medicinal plants, organ formation and development are controlled by complex interactions among genetic and environmental factors. The transcriptome data in publicly available libraries are insufficient and limited to describe the complex mechanisms of gene expression, as well as the genetic characteristics of species. Therefore, new generation of high-throughput sequencing technologies has been used as a powerful and cost-efficient tool for research on non-model organisms (Gomes-Carneiro et al. 1998; Haralampidis et al. 2015). According to WHO, medicinal plants play a vital role in pharmaceutically industry-relevant secondary metabolites, in which *Tribulus terrestris* is an important traditional medicinal herb with numerous pharmaceutical activities. However, there has been some research on genomics, but no research on the transcriptome of *T. terrestris*. This research paper mainly focuses on the transcriptome of *T. terrestris* and reveals diosgenin biosynthetic genes. We described a comparative transcriptome analysis of *T. terrestris* (root, fruit, and leaf tissues). The dataset provided here is useful in understanding the diosgenin biosynthesis pathway.

Through Illumina, ∼7.9 GB clean data were generated from RNA-seq libraries in replicate form whole parts of *T. terrestris*. Sample-wise, 482 (4,823,004) million paired-end reads were collected and these are reliable for *de novo* transcriptome characterization and precise gene expression pattern quantification (Zhou et al. 2019). 16889795 (41.10%) duplicate filtered reads assembled into 200471 unigenes with an average N50 length of 956 bp, which is similar to that of previously reported non-model herbs and plants, *Trigonella foenum-graecum, Trillium govanianum*, and *Dioscorea zingiberensis* (Petr et al. 2014). The results are comparable with the obtained unigenes in the recently published transcriptome analyses of other plant species, such as *H. ammodendron* (N50 = 1,354 bp, average length = 728 bp) (Haralampidis et al. 2002), *Reaumuria soongorica* (N50 = 1,109 bp, average length = 677 bp) (Abe et al. 1993), and radish (*Raphanus sativus* L.) (N50 = 1,256 bp, average length = 820 bp) (Xu et al. 2004). Longer unigenes may be obtained because of the developed Trinity software, which is a powerful software package for *de novo* assembly and generates increased number of full-length transcripts (Perez-Labrada et al. 2011). GC content 40251760 (43.34%) may be attributed to the ability of *T. terrestris* to adapt in extreme temperature as GC content play important role in gene regulation, physical description of genome and nucleic acid stability (Ge et al. 2014), besides reflecting high quality sequencing run. While, G+C percentage of *T. terrestris* (43.34%) was higher than *Arabidopsis* (42.5%) (Goossens et al. 2015). Despite being a non-model herb of *T. terrestris* transcripts were annotated with multiple public databases successfully assigned putative functions to over 100753 of transcripts. Nonetheless, 11429 transcripts could not be annotated possibly belongs to the un-translated regions or represents the species-specific gene-pool (Roy et al. 2021; Liu et al. 2015). The best match for each unigenes search against the Nr and KEGG databases was of help to assign GO functional annotation under biological process, cellular component, and molecular function categories.

The assignment of GO keywords to a substantial number of transcripts in *T. terrestris* suggests the presence of several gene families. The mapping of transcripts to the KEGG database in this work identified all genes involved in the steroidal diosgenin pathway, assisting in the understanding of the biological function and interaction of genes involved in primary and secondary metabolites. A total of 37292 transcripts were annotated and classified using the COG categorization approach into 25 functional categories. It has long been recognized that transcription factors (TFs) play a critical role in gene expression regulation by binding to the promoters of one or many genes. Our data indicate that the transcription factor family’s bHLH, NAC, MYB related family, AP2/ERF, C2H2, WRKY, C3H, MYB, Bzip, and B3 are highly expressed and essential to the regulation of secondary metabolites in plants. TSAR1 and TSAR2 are members of the bHLH family and regulate triterpene diosgenin production in *Medicago truncatula* (Jiang et al. 2015). As a result, the TFs identified in this study might be studied as possible regulators of *T. terrestris* steroidal diosgenin production. On the other hand, RNA-seq is incapable of identifying genes or transcripts with low expression levels, which is particularly critical for regulatory genes (Graveley et al. 2011; Malone et al. 2011). In completely sequenced modENCODE embryo samples with low expression, the transcription factors (TFs) involved in sexual dimorphism in flies, doublesex (*dsx*) and fruitless (*fru*), cannot be detected by RNA-Seq (Loven et al. 2012). These genes will have an effect on both the estimation of transcript expression and functional research.

Gene expression studies are now widely used in a variety of fields to find possible regulators of complex biochemical pathways by comparing transcriptional levels across tissues and developmental stages (Kim et al. 2014). A substantial number of DEGs transcripts were found in pairwise alignments of *T. terrestris* root, fruit, and leaf. All unigenes encoding enzymes involved in the glycolytic, mevalonate (MVA), MEP/DOX, mono-terpenoids, and steroidal pathways for diosgenin biosynthesis from acetyl-CoA to squalene were found to support the annotation. In transcriptomic analysis, encoding sequences related with the *Glycolotic* pathway including genes *HMGS* (EC: 2.3.3.10, 6 transcripts), *HMGR* (EC: 1.1.1.34, 24 transcripts); *mevalonate* pathway *MVK (EC:* 2.7.1.36, 1 transcripts*), PMK* (EC: 2.7.4.2, 4 transcripts), *MVD* (EC: 4.1.1.33, 4 transcripts); *MEP/DOX* pathway *DXD* (EC:2.2.1.7, 1 transcripts), *DXR* (EC:1.1.1.267, 1 transcripts); *mono-terpenoids* pathway *IDI* (EC: 5.3.3.2, 4 transcripts), *steroids* pathway *GPPS* (EC: 2.5.1.29, 6 transcripts), *SQS* (EC: 1.14.14.17, 12 transcripts), *SMT1* (EC: 2.1.141, 3 transcripts), *DWF7* (EC:1.3.1.21, 5 transcripts), and *FK* (EC:1.3.1.70, 5 transcripts) of *T. terrestris* represented the highest number of unigenes involved in the synthesis of phytosterols, carotenoids, gibberellins, triterpenoid and steroidal saponins Supplementary Table S3. The highest expression of MEP and *steroids* pathway genes were identified during this study and also reported in earlier studies (Goossens et al. 2015). The cyclization of 2, 3-oxidosqualene is a branch point of diosgenin synthesis. A total of 6 transcripts were annotated of housekeeping genes (alpha-tubulin) of our transcriptome. The high expression of *alpha-tubulin* synthase in leaf of *T. terrestris*, an important enzyme related to triterpenoid diosgenin biosynthesis at later stages. Similar tissue-specific concentrations of triterpenoid diosgenin have already been reported in other herbs (Petr et al. 2014). Moreover, *alpha-tubulin, HMGR, DXR, IDI, GPPS, MVD, SQE, DWF7, HMGR*, and *DXR* genes were up-regulated in which, MVD gene in leaf was down-regulated. While, *MVK* gene was absent in all parts. Further characterization of these candidate enzymes is needed to confirm the pathway of triterpenoid diosgenin biosynthesis in *T. terrestris*.

In the analysis of the SSR polymorphism loci of the *T. terrestris* transcriptome with 150bp length and 245069 unigenes by using the MISA software, and 17623 transcriptomic sequences were generated. A total of 15551 SSR loci were detected with a frequency and an average distribution distance of 179930901 bp. In the *T. terrestris* SSR loci, the most frequent repeat type is mono-nucleotide with 3,394 (48.27%), followed by di-, tri-, tetra-, penta-nucleotide repeats, with 9292, 6580, 146, and 4 respectively. This distribution frequency differs from those of most plant genomes, such as field *Pisum sativum, Vicia faba*, and *Medicago sativa*, in which the most abundant repeat motif is tri-nucleotide (57.7%, 61.7%, and 61.19%, respectively) (Sukhjiwan et al. 2012; Liu et al. 2013); in *Sesamum indicum*, the most abundant repeat motif is di-nucleotide repeat motifs (Wei et al. 2011). *Medicago sativa, Pisum sativum, Vicia faba*, and *Polygoni cuspidatum* do not have mononucleotide repeat sequences, which could be due to the different standards used in SSR search criteria (Liu et al. 2013). In this study, we explored the mono-nucleotide repeat motifs in *T. terrestris*; during the process, a condition where the mono-nucleotide repeat is dominant was generated, which decreases the number of other nucleotide repeats.

## 5. Conclusions

Pharmaceutical industries depend primarily on medicinal plants as a source of botanical raw materials. In this study, we performed whole transcriptome analysis of *T. terrestris* root, fruit, and leaf tissues. All of the key genes involved in the diosgenin biosynthesis pathway can be studied futuristically for bioactive molecule up-scaling. The highest expression of important pathway genes and regulatory candidates in *T. terrestris* is reported in the root and leaf, implying that these are the sites of steroidal diosgenin synthesis. The current study findings will lay the groundwork for future multi-omics studies in *T. terrestris* and related species in order to gain a better understanding of steroidal diosgenin synthesis and accumulation. The data collected will aid in the discovery of new genes and functional genomics in *T. terrestris* for the purpose of genetic improvement and conservation studies.

## Acknowledgments

We are thankful to the Director and Head of Department of Botany, Dayalbagh Educational Institute, Dayalbagh (Agra).

## Supplementary information

### Supplementary figures

**Supplementary Figure S1:** ATGC composition of root, fruit, and leaf tissues *T. terrestris* transcripts.

**Supplementary Figure S2:** Length of all assembled transcripts (minimum (200-300) and maximum (5000) bp) of *T. terrestris*.

**Supplementary Figure S3:** GC contain of fastaqc file (R1 and R2) of *T. terrestris*

**Supplementary Figure S4:** Functional annotation: Venn diagram showing functional annotation details of transcripts with NR, TAIR10, Swiss-Prot and COG databases of *T. terrestris* tissues.

**Supplementary Figure S5:** Homologous Similarity of *T. terrestris*: Similarity of unigenes annotated using the Nr database. E-value distribution of best BLAST hits for each unigene (E-value <1e-6), similarity distribution of top BLAST hits for each unigene, and distribution of the most homologous sequence results for each unigene by species (E-value <1e-6).

**Supplementary Figure S6:** Top 20 Pfam domains in *T. terrestris*.

**Supplementary Figure S7:** Enzymes are assembled in diosgenin pathway analysis *T. terrestris*.

**Supplementary Figure S8:** Function annotation classifications: Classification of transcripts into major transcription factor (TF) families of *T. terrestris*.

**Supplementary Figure S9:** Phylogenetic analysis of all diosgenin related genes transcripts of root, fruit, and leaf tissues of *T. terrestris*. The distances between each transcript were constructed by neighbor-joining method using *MEGAX* software.

## Supplementary tables

**Supplementary Table S1:** Transcriptome distribution of CDS with the BLASTX of all parts of *Tribulus terrestris*

**Supplementary Table S2:** *De novo* transcriptome analysis of unigenes of KEGG mapping pathways analysis of root, fruit, and leaf tissues of *T. terrestris*.

**Supplementary Table S3:** List of genes which are related to diosgenin in *T. terrestris*

**Supplementary Table S4:** Primer Identification of internal control genes and some of these genes are used for validation of qRT-PCR analysis of *T. terrestris*

**Supplementary Table S5:** Transcriptome data analysis SSRs or microsatellite data of *T. terrestris*

## References

1. Abe I, Rohmer M, Prestwich GD (1993) Enzymatic cyclization of squalene and oxidosqualene to sterols and triterpenes. Chem Rev. 3, 2189–206. https://doi.org/10.1021/cr00022a009.

2. Akram M, Asif HM, Akhtar N, Shah PA, Uzair MU, Shaheen G, Shamim T, Shah SA (2011) Tribulus terrestris Linn.: a review article. Journal of Medicinal Plants Research 18:5(16). https://doi.org/10.5897/JMPR.9001271.

3. Altschul SF, Gish W, Miller W, Myers EW, Lipman DJ (1990) Basic local alignment search tool. J. Mol. Biol 215, 403–10. https://doi.org/10.1016/S0022-2836(05)80360-2.

4. Amos B, Rolf A (2003) The SWISS-PROT protein sequence database and its supplement TrEMBL in 2000 Nucleic Acids Res. 28, 45–48. https://doi:10.1093/nar/28.1.45

5. Audic S, Claverie J-M (1997) The significance of digital gene expression profiles. Genome Research 7: 986–995. https://doi:10.1101/gr.7.10.986.

6. Bairoch B, Apweiler R (2000) The SWISS-PROT protein sequence database and its supplement TrEMBL, Nucleic Acids Res 28: 45–48. https://doi.org/10.1093/nar/28.1.45

7. Bolger AM, Lohse M, Usadel B (2014) Trimmomatic: a flexible trimmer for Illumina sequence data. Bioinformatics. 30, 2114–20. https://doi.org/10.1093/bioinformatics/btu170.

8. Chen Y, Shi C, Huang Z, Zhang Y, Li S, Li Y, Ye J, Yu C, Li Z (2018) SOA Pnuke: a Map Reduce acceleration supported software for integrated quality control and preprocessing of high-throughput sequencing data. Gigascience. 7:1–6.

9. Chhatre S, Nesari T, Somani G, Kanchan D, Sathaye S (2014) Phytopharmacological overview of Tribulus terrestris Pharmacogn Rev. 45–51.

10. Christenhusz MM, Byng JW (2016) The number of known plants species in the world and its annual increase. Phytotaxa. Magnolia Press. 261 (3): 201–217. doi:10.11646/phytotaxa.261.3.1.

11. Crawford JE, Guelbeogo WM, Sanou A, Traore A, Vernick KD, Sagnon NF, Lazzaro BP (2010) De novo transcriptome sequencing in Anopheles funestus using Illumina RNA-Seq technology. PLoS One. 5, e14202.

12. Finn RD, Bateman A, Clements J, Coggill P, Eberhardt RY, Eddy SR, Heger A, Hetherington K, Holm L, Mistry J (2014) Pfam: the protein families database. Nucleic Acids Res. 42, D222–30.

13. Fujita M, Fujita Y, Noutoshi Y, Takahashi F, Narusaka Y, Yamaguchi-Shinozaki K, Shinozaki K (2006) Crosstalk between abiotic and biotic stress responses: a current view from the points of convergence in the stress signaling networks. Current opinion in plant biology. 9(4), 436–442.

14. Ge X, Chen H, Wang H, Shi A, Liu K (2014) De novo assembly and annotation of Salvia splendens transcriptome using the Illumina platform. PloS one 9(3), e87693.

15. Gene Ontology (2004) Consortium, The Gene Ontology (GO) database and informatics resource. Nucleic Acids Research. 32, 258–261.

16. Gomes-Carneiro MR, Felzenszwalb I, Paumgartten FJ (1998) Mutagenicity testing of (±)-camphor, 1,8-cineole, citral, citronellal, (-) menthol and terpineol with the Salmonella/microsome assay. Mutat. Res. - Genet. Toxicol. Environ. Mutagen 416, 129–136. https://doi.org/10.1016/S1383-5718(98)00077-1.

17. Goossens A (2015) The bHLH Transcription Factors TSAR1 and TSAR2 Regulate Triterpene Saponin Biosynthesis in Medicago truncatula. Plant physiology. pp-01645.

18. Grabherr MG, Brian J, et al (2013) Friedman, Trinity: reconstructing a full-length transcriptome without a genome from RNA-Seq data. Nat. Biotechnol 29, 644–52.

19. Graveley BR, Brooks AN, Carlson JW, Duff MO, Landolin JM, Yang L, Artieri CG, Baren MJ, Boley N, Booth BW (2011) The developmental transcriptome of Drosophila melanogaster. Nature 471:473–9.

20. Haas JB, Papanicolaou A, Yassour M, Grabherr M, Friedman N, Regev A (2014) De novo transcript sequence reconstruction from RNA-Seq: reference generation and analysis with Trinity HHS Public Access 8(8): 10–1038.

21. Haas JB, PapanicolaouA, Yassour M, Grabherr M, Friedman N, Regev A (2014) De novo transcript sequence reconstruction from RNA-Seq: reference generation and analysis with Trinity HHS Public Access 8(8), 10–1038.

22. Haralampidis K, Trojanowska M, Osbourn AE (2002) Biosynthesis of triterpenoid saponins in plants. Adv Biochem Eng Biotechnol. 75, 31–49.

23. He M, Wang Y, Hua W, Zhang Y, Wang Z (2012) De Novo Sequencing of Hypericum perforatum Transcriptome to Identify Potential Genes Involved in the Biosynthesis of Active Metabolites. PLoS ONE. 7 doi: 10.1371/journal.pone.0042081.

24. Hua W, Zhang Y, Song J, Zhao L, Wang Z (2011) De novo transcriptome sequencing in Salvia miltiorrhiza to identify genes involved in the biosynthesis of active ingredients. Genomics. 98:272–279.

25. Jiang Z, Zhou X, Li R, Michal JJ, Zhang S, Dodson MV, Zhang Z, Harland RM (2015) Whole transcriptome analysis with sequencing: methods, challenges and potential solutions. Cellular and molecular life sciences: CMLS., 72(18):, 3425–39.

26. Kanehisa M, Goto S, Sato Y, Kawashima M, Furumichi M (2014) Tanabe Data, information, knowledge and principle: Back to metabolism in KEGG. Nucleic Acids Res 42, D199–205. https://doi.org/10.1093/nar/gkt1076.

27. Kelemen O, Convertini P, Zhang Z, Wen Y, Shen M, Falaleeva M, Stamm S (2013) Function of alternative splicing, Gene, 514, pp. 1–30.

28. Kim JA, Roy NS, Lee I, Choi AY, Choi BS, Yu YS, Park N, Park KC, Kim S, Yang H, Choi IY (2019) Genome-wide transcriptome profiling of the medicinal plant Zanthoxylum planispinum using a single-molecule direct RNA sequencing approach, Genomics, Pages 973–979.

29. Kim WS, Haj-Ahmas Y (2014) Evaluation of Plant RNA Integrity Number (RIN) generated using an Agilent BioAnalyzer 210. Norgen Biotek Corp.

30. Kumar S, Kalra S, Singh B, Kumar A, Kaur J, Singh K (2016) RNA-Seq mediated root transcriptome analysis of Chlorophytum borivilianum for identification of genes involved in saponin biosynthesis. Funct. Integr. Genom.;16:37–55. doi: 10.1007/s10142-015-0465-9.

31. Lehnert EM, Mouchka ME, Burriesci MS, Gallo ND, Schwarz JA, Pringle JR (2014) Extensive Differences in Gene Expression Between Symbiotic and Aposymbiotic Cnidarians, G3 Genes|Genomes|Genetics, 4:(2), 277–295, https://doi.org/10.1534/g3.113.009084.

32. Lindgreen S (2012) Adapter Removal: easy cleaning of next-generation sequencing reads. BMC Res Notes. 5–337.

33. Lindgreen S (2012) Adapter Removal: easy cleaning of next-generation sequencing reads. BMC Res Notes.5–337.

34. Liu Y, Zhang P, Song M, Hou J, Qing M, Wang W, et al (2015) Transcriptome Analysis and Development of SSR Molecular Markers in Glycyrrhiza uralensis Fisch. PLoS ONE 10(11): e0143017. https://doi.org/10.1371/journal.pone.0143017.

35. Liu ZP, Chen TL, Ma LC, Zhao ZG, Zhao Patrick X, Nan ZB, et al (2013) Global Transcriptome Sequencing using the Illumina Platform and the Development of EST-SSR Markers in Autotetraploid Alfalfa. PLOS One.; 8(12): e83549. pmid:24349529.

36. Livak KL, Schmittgen TD (2001) Analysis of relative gene expression data using real-time quantitative PCR and the 2(-Delta Delta C(T)) Method. Methods., 25(4), 402–8.

37. Loven J (2012) Revisiting global gene expression analysis. Cell. 151(3): 476–482.

38. Lulin H, Xiao Y, Pei S, Wen T, Shangqin H (2012) The First Illumina-Based De Novo Transcriptome Sequencing and Analysis of Safflower Flowers. PLoSONE 7(6); e38653. doi:10.1371/journal.pone.0038653.

39. Malone JH, Oliver B (2011) Microarrays, deep sequencing and the true measure of the transcriptome. BMC Biol. 9:34.

40. Manfred G, Grabherr BJ (2013) Trinity: reconstructing a full-length transcriptome without a genome from RNA-Seq data, Nat Biotechnol. 29, 644–652. https://doi:10.1038/nbt.1883.

41. Mehrafarin A, Qaderi A, Rezazadeh S, Naghdi Badi H, Noormohammadi G, Zand E (2010) Bioengineering of Important Secondary Metabolites and Metabolic Pathways in Fenugreek (Trigonella foenum-graecum L.) J. Med. Plants.;9:1–18.

42. Miraj S (2016) Tribulus terrestris: Chemistry and pharmacological properties Der Pharma Chemica. 142–147.

43. Mistry J, Finn RD, Eddy SR, Bateman A, Punta M (2013) Challenges in homology search: HMMER3 and convergent evolution of coiled-coil regions. Nucl Acids Res., 41(12):e121.

44. Moraortiz M, Swain MT, Vickers MJ, Hegarty MJ, Kelly R, Smith LJ, Skot L (2016) De novo transcriptome assembly for gene identification, analysis, annotation, and molecular marker discovery in Onobrychis viciifolia. BMC Genomics. 17 (1). 756. ISSN 1471-2164 doi:https://doi.org/10.1186/s12864-016-3083-6

45. Moriya Y, Itoh M, Kanehisa, Okuda S, Yoshizawa AC, Kanehisa M (2007) KAAS: An automatic genome annotation and pathway reconstruction server. Nucleic Acids Res W182–W185, https://doi.org/10.1093/nar/gkm321.

46. Muranaka T, Saito K (2013) Phytochemical genomics on the way, Plant Cell Physiol., 54, pp. 645–646.

47. Nakasugi K, Kenlee N, Ross C, Julia B, Peter W (2014) Combining Transcriptome Assemblies from Multiple De Novo Assemblers in the Allo-Tetraploid Plant Nicotiana benthamiana, PLoS One. 9(3):, e91776. doi: 10.1371/journal.pone.0091776.

48. Patra B, Schluttenhofer C, Wu Y, Pattanaik S, Yuan L (2013) Transcriptional regulation of secondary metabolite biosynthesis in plants. Biochimica et BiophysicaActa (BBA)-Gene Regulatory Mechanisms. 1829(11), 1236–1247.

49. Perez-Labrada K (2011) ‘Click’ synthesis of triazole-based spirostan saponin analogs. Tetrahedron. 67(40), 7713–7727.

50. Perteal G, Huang XQ, Liang F, Antonescu V, Sultanal R, Karamycheval S (2003) TIGR gene indices clustering tools (TGICL): a software system for fast clustering of large EST datasets. Bioinformatics. 19, 651–2.

51. Petr S (2014) Ecological and evolutionary significance of genomic GC content diversity in monocots. Proceedings of the National Academy of Sciences. 111(39), E4096–E4102.

52. Qureshi A, Declan P, Naughton DP, Petroczi A (2014) A Systematic Review on the Herbal Extract Tribulus terrestris and the Roots of its Putative Aphrodisiac and Performance Enhancing Effect. Journal of Dietary Supplements. 64–79.

53. Roy NS, Choi IY, Um T, Jeon MJ, Kim BY, Kim YD, Yu JK, Kim S, Kim NS (2021) Gene Expression and Isoform Identification of PacBio Full-Length cDNA Sequences for Berberine Biosynthesis in Berberis koreana; Plants, 10, 1314. https://doi.org/10.3390/plants10071314.

54. Saito K (2013) Phytochemical genomics-a new trend, Curr. Opin. Plant Biol., 16, pp. 373–380.

55. Schluttenhofer C, Yuan L (2015) Regulation of Specialized Metabolism by WRKY Transcription Factors. Plant Physiol., 167: 295–306. doi: 10.1104/pp.114.251769.

56. Sonawane PD, Pollier J, Panda S, Szymanski J, Massalha H, Yona M, Unger T, Malitsky S, Arendt P, Pauwels L, et al. (2016) Plant cholesterol biosynthetic pathway overlaps with phytosterol metabolism. Nat. Plants.;3 doi: 10.1038/nplants.2016.205.

57. Storey JD, Tibshirani R (2003) Statistical significance for genomewide studies. Proc. Natl. Acad. Sci. USA., 100, 9440–9445, doi: 10.1073/pnas.1530509100.

58. Sukhjiwan K, Pembleton LW, Cogan Noel Ol, Savin Keith W, Tony L, Jeffrey P, et al. (2012) Transcriptome sequencing of field pea and faba bean for discovery and validation Of SSR genetic marker. BMC Genomics.; 13: 104. pmid:22433453.

59. Trapnell A, Williams BA, Pertea G, Mortazavi A, Kwan G, Baren MJ, Salzberg SL, Wold BJ, Pachter L (2010) Transcript assembly and quantification by RNA-Seq reveals unannotated transcripts and isoform switching during cell differentiation. Nat. Biotechnol. 28:511–515. doi: 10.1038/nbt.1621.

60. Tyagi P, Ranjan R (2021) Comparative study of the pharmacological, phytochemical and biotechnological aspects of Tribulus terrestris Linn. and Pedalium murex Linn: An overview. Acta Ecologica Sinic, https://doi.org/10.1016/j.chnaes.2021.07.008.

61. Unamba CI, Nag A, Sharma RR (2015) Next Generation Sequencing technologies: The doorway to the unexplored genomics of non-model plants. Frontiers in plant science 6.

62. Vasait, Rajendrabhai C (2017) Detection of Phytochemical and Pharmacological Properties of Crude Extracts of Tribulus terrestris Collected from Tribal Regions of Baglan (M.S.) India. International Journal of Pharmacognosy and Phytochemical Research. 508–511.

63. Volkman JK (2005) Sterols and other triterpenoids: Source specificity and evolution of biosynthetic pathways. Org. Geochem.;36:139–159. doi: 10.1016/j.orggeochem.2004.06.013.

64. Vranova E, Coman D, Gruissem W (2013) Network analysis of the MVA and MEP pathways for isoprenoid synthesis. Annu. Rev. Plant Biol. 2013;64:665–700. doi: 10.1146/annurev-arplant-050312-120116.

65. Wang S, Wang B, Hua W, Niu J, Dang K., Qiang, Y, Wang Z (2017) De novo assembly and analysis of Polygonatum sibiricum transcriptome and identification of genes involved in polysaccharide biosynthesis. Int J Mol Sci. 18.

66. Wang Z, Gerstein M, Snyder M (2009) RNA-Seq: a revolutionary tool for transcriptomics, Nat. Rev. Genet., 10, pp. 57–63.

67. Wei WL, Qi XQ, Wang LiH, Zhang YX, Hua W, Li DH, et al. (2011) Characterization of the sesame (Sesamum indicum L.) global transcriptome using Illumina paired-end sequencing and development of EST-SSR markers. BMC Genomics.; 12: 451. pmid:21929789.

68. Xu R, Fazio GC, Matsuda SP (2004) On the origins of triterpenoid skeletal diversity. Phytochemistry. 65, 261–91.

69. Yin J, Li X, Zhan Y, Li Y, Qu Z, Sun L, Wang S, Yang J, Xiao J (2017) Cloning and expression of BpMYC4 and BpbHLH9 genes and the role of BpbHLH9 in triterpenoid synthesis in birch. BMC Plant Biol., 17 doi: 10.1186/s12870-017-1150-z.

70. Paul P, Singh SK, Patra B, Sui X, Pattanaik S, Yuan L (2017) A differentially regulated AP2/ERF transcription factor gene cluster acts downstream of a MAP kinase cascade to modulate terpenoid indole alkaloid biosynthesis in Catharanthus roseus. New Phytol., 213:1107–1123. doi: 10.1111/nph.14252.

71. Zhou C, Xiaohua L, Zhou Z, Li C, Zhang Y (2019) Comparative Transcriptome Analysis Identifies Genes Involved in Diosgenin Biosynthesis in Trigonella foenum-graecum L.; Molecules, 24(1): 140. doi: 10.3390/molecules24010140.

