## Supplemental table and figures for "Illumina-based Whole *De Novo* Transcriptome Profiling and Diosgenin Biosynthetic pathway of *Tribulus terrestris:* A Medicinal herb": SUPPLIMENTARY FILE.docx

**Supplementary information**

**Supplementary Figure S1:** ATGC composition of root, fruit, and leaf tissues *T. terrestris* transcripts.

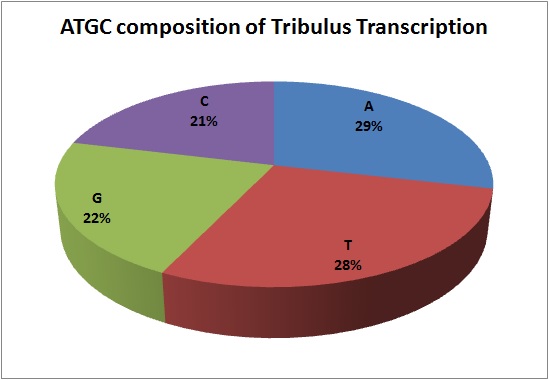

**Supplementary Figure S2:** Length of all assembled transcripts (minimum (200-300) and maximum (5000) bp) of *T. terrestris.*

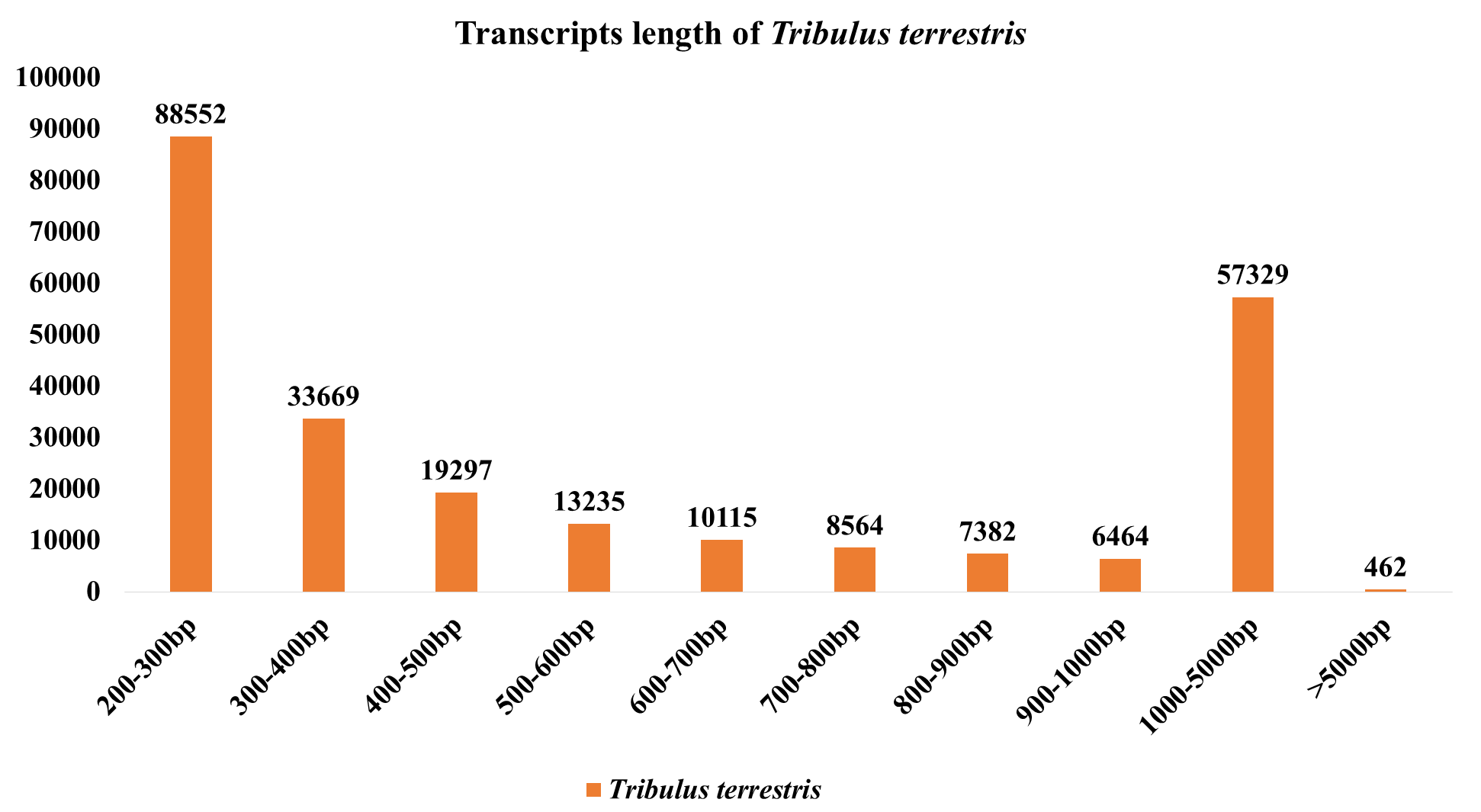

**Supplementary Figure S3:** GC contain of fastaqc file (R1 and R2) of *T. terrestris*

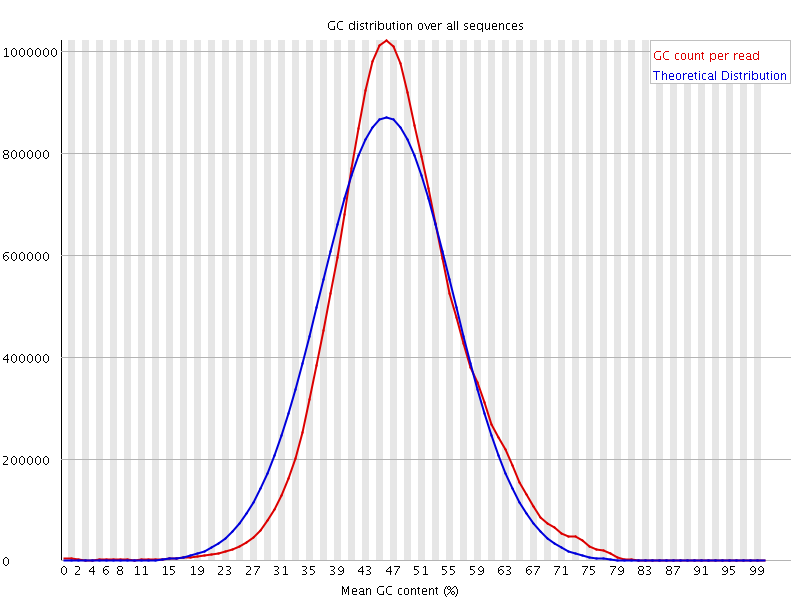

*T. terrestris fastaqc* R1

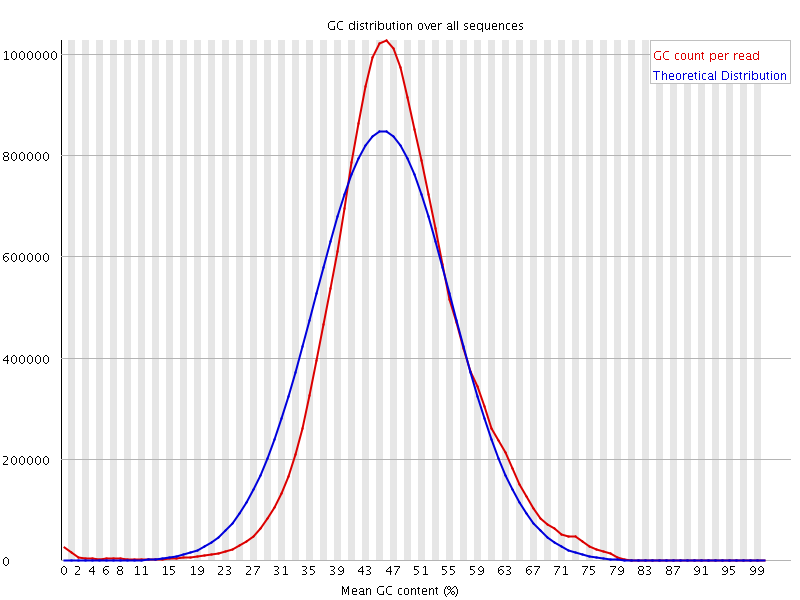

*T. terrestris fastaqc* R2

**Supplementary Figure S4:** Functional annotation: Venn diagram showing functional annotation of all common unigenes details of with COG, Uniprot, Pfam, and KEGG databases of *T. terrestris* tissues.

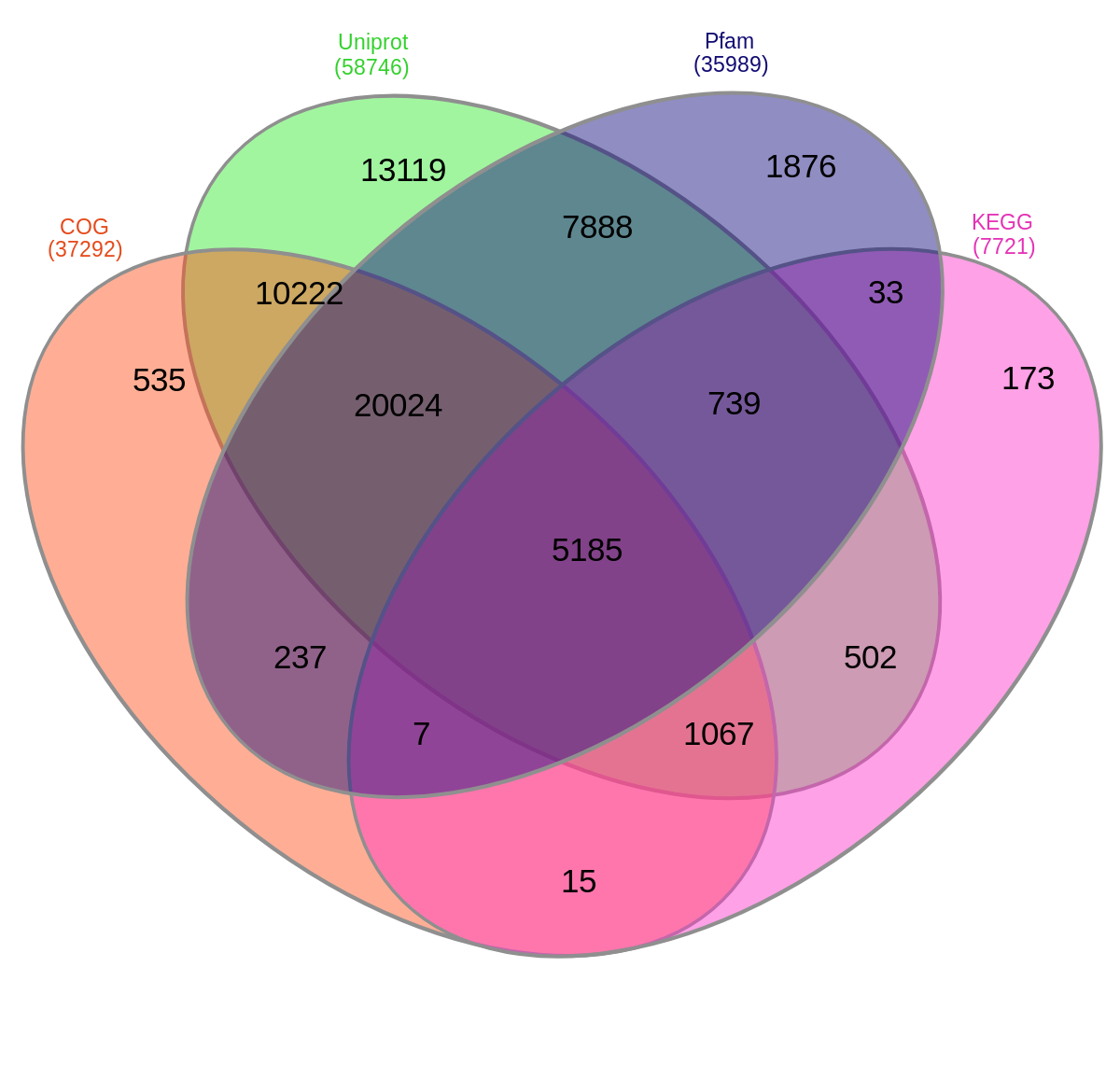

**Supplementary Figure S5:** Homologous similarity of *T. terrestris*: Similarity of unigenes annotated using the Nr database. E-value distribution of best BLAST hits for each unigene (E-value <1e-6), similarity distribution of top BLAST hits for each unigene, and distribution of the most homologous sequence results for each unigene by species (E-value <1e-6).

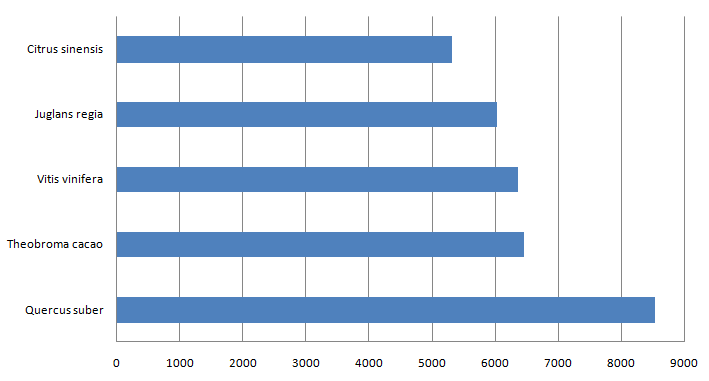

**Supplementary Figure S6:** Top 20 Pfam domains in *T. terrestris,*

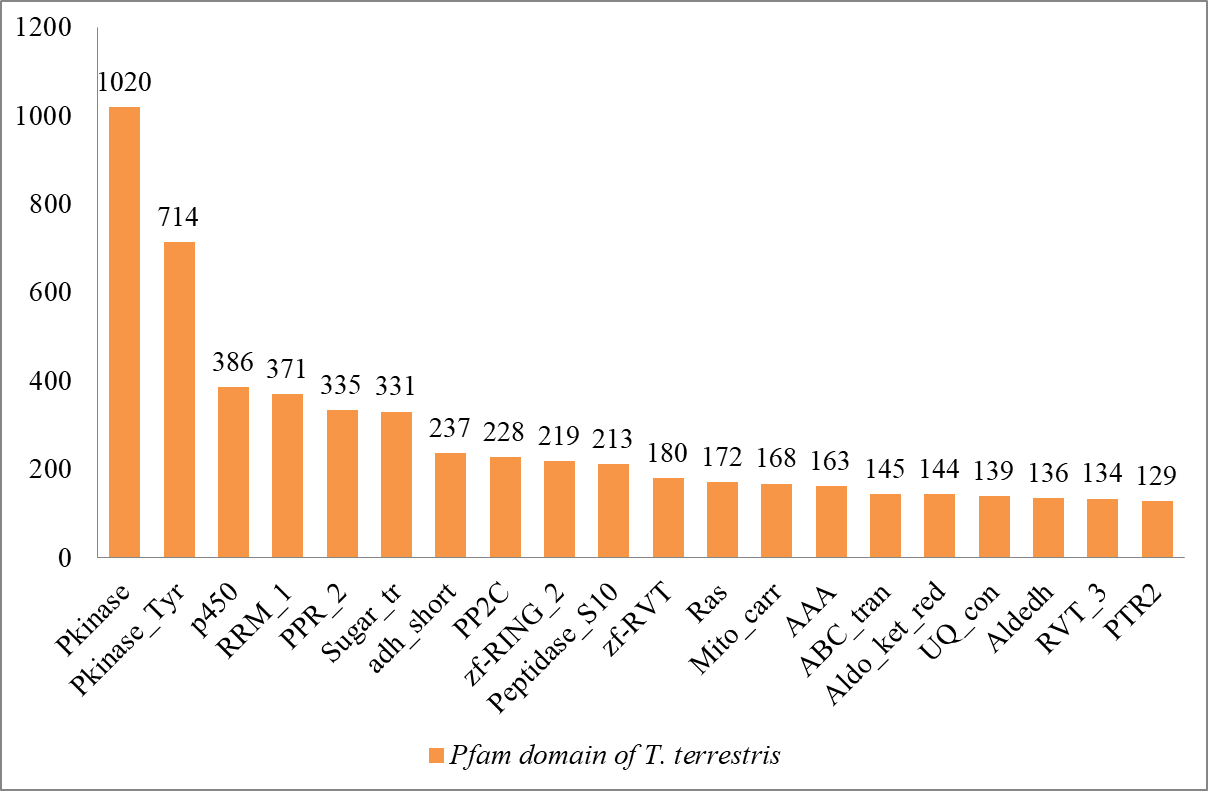

**Y axis- No. of unigenes**

**X axis- No. of enzymes**

**Supplementary Figure S7:** Enzymes are assembled in diosgenin pathway analysis *T. terrestris.*

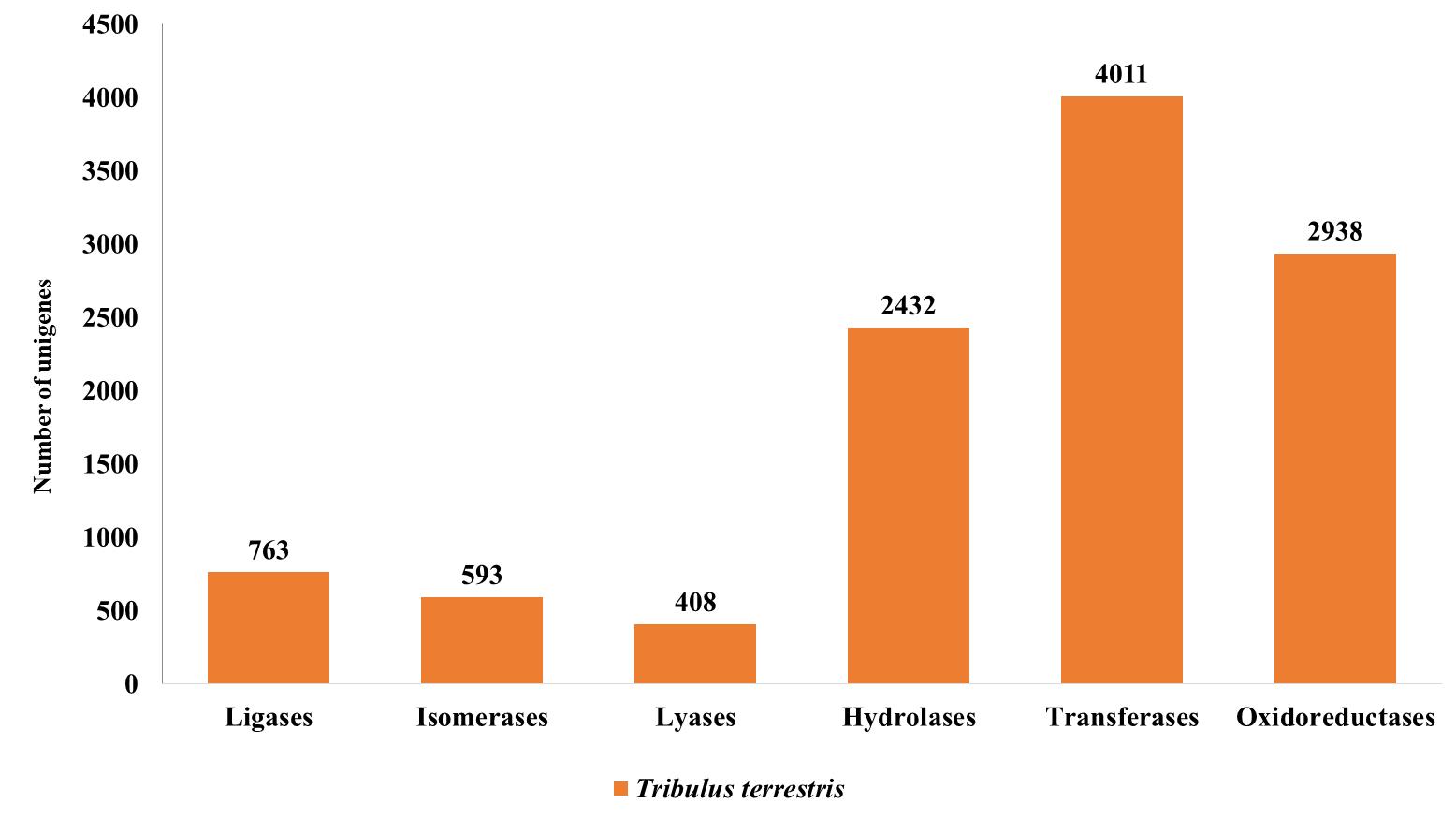

**Supplementary Figure S8:** Function annotation classifications: Classification of transcripts into major transcription factor (TF) families of *T. terrestris.*

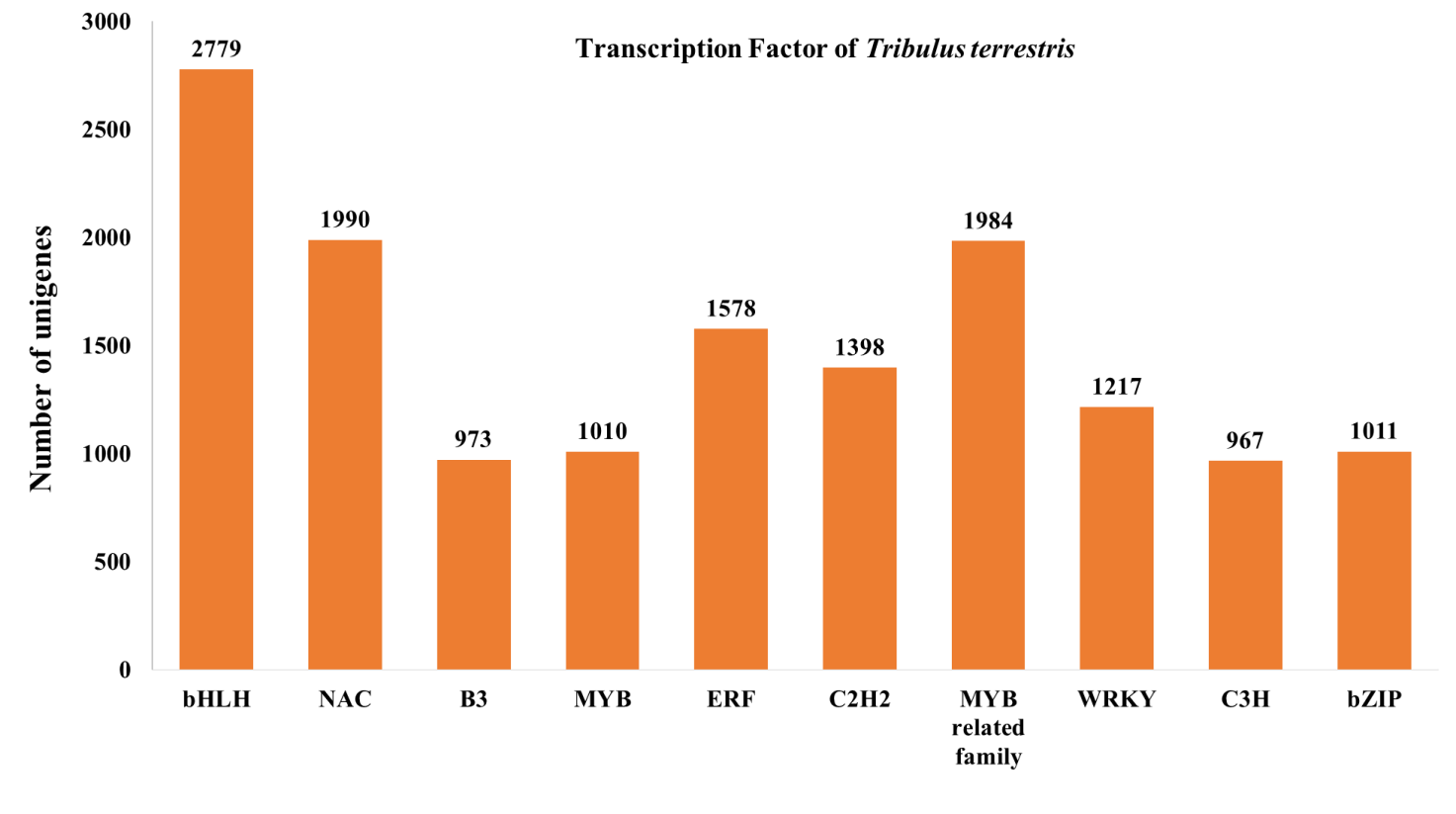

**Supplementary Figure S9:** Phylogenetic analysis of all diosgenin related genes transcripts of root, fruit, and leaf tissues of *T. terrestris.* The distances between each transcript were constructed by neighbor-joining method using *MEGAX* software.

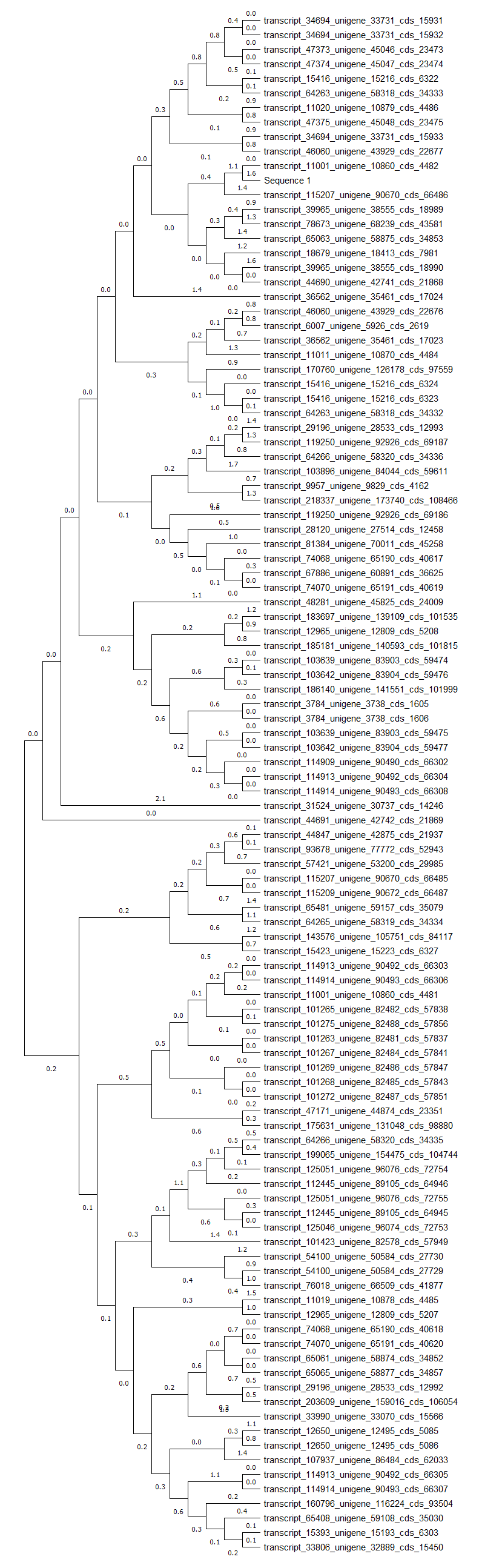

**Supplementary Table S1:** Transcriptome distribution of CDS with the BLASTX of all parts of *Tribulus terrestris*

| Samples | No. of CDS | No. of CDS with BLAST hits | No. of CDS without BLAST hits | GO distribution of BLAST hits | | |
| --- | --- | --- | --- | --- | --- | --- |
|  |  |  |  | **Biological processes** | **Molecular functions** | **Cellular compounds** |
| Root, fruit, and leaf of *T. terrestris* | **112182** | **100753** | **11429** | **33258** | **37044** | **26597** |

**Supplementary Table S2:** *De novo* transcriptome analysis of unigenes of KEGG mapping pathways analysis of root, fruit, and leaf tissues of *T. terrestris.*

| **Pathways analysis** | **Pathways categorises** | **Root, fruit, and leaf of *T. terrestris*** |
| --- | --- | --- |
| 09100 Metabolism | (09101)- Carbohydrate metabolism | 808 |
|  | (09102)- Energy metabolism | 457 |
|  | (09103)- Lipid metabolism | 406 |
|  | (09104)- Nucleotide metabolism | 291 |
|  | (09105)- Amino acid metabolism | 535 |
|  | (09106)- Metabolism of other amino acids | 182 |
|  | (09107)- Glycan biosynthesis and metabolism | 178 |
|  | (09108)- Metabolism of cofactors and vitamins | 256 |
|  | (09109)- Metabolism of terpenoids and polyketides | 126 |
|  | (09110)- Biosynthesis of other secondary metabolites | 182 |
|  | (09111)- Xenobiotics biodegradation and metabolism | 146 |
| 09120 Genetic Information  Processing | (09121)- Transcription | 338 |
|  | (09122)- Translation | 783 |
|  | (09123)- Folding, sorting and degradation | 539 |
|  | (09124)- Replication and repair | 371 |
| 09130 Environmental  Information Processing | (09131)- Membrane transport | 30 |
|  | (09132)- Signal transduction | 1165 |
|  | (09133)- Signalling molecules and interaction | 0 |
| 09140 Cellular Processes | (09141)- Transport and catabolism | 602 |
|  | (09142)- Cell motility | 45 |
|  | (09143)- Cell growth and death | 578 |
|  | (09144)- Cellular community - Eukaryotes | 79 |
|  | (09145)- Cellular community - Prokaryotes | 66 |
| 09150 Organismal  Systems | (09149)- Aging | 118 |
|  | (09151)- Immune system | 342 |
|  | (09152)- Endocrine system | 375 |
|  | (09153)- Circulatory system | 76 |
|  | (09154)- Digestive system | 84 |
|  | (09155)- Excretory system | 68 |
|  | (09156)- Nervous system | 269 |
|  | (09157)- Sensory system | 30 |
|  | (09158)- Development | 59 |
|  | (09159)- Environmental adaptation | 351 |
| 09160 Human Diseases | (09161)- Cancers: Overview | 353 |
|  | (09162)- Cancers: Specific types | 238 |
|  | (09163)- Immune diseases | 70 |
|  | (09164)- Neurodegenerative diseases | 346 |
|  | (09165)- Substance dependence | 96 |
|  | (09166)- Cardiovascular diseases | 85 |
|  | (09167)- Endocrine and metabolic diseases | 151 |
|  | (09171)- Infectious diseases: Bacterial | 261 |
|  | (09172)- Infectious diseases: Viral | 687 |
|  | (09174)- Infectious diseases: Parasitic | 81 |
|  | (09175)- Drug resistance: Antimicrobial | 3 |
|  | (09176)- Drug resistance: Antineoplastic | 81 |
| 09180 Brite Hierarchies | (09181)- Protein families: Metabolism | 1243 |
|  | (09182)- Protein families: Genetic information processing | 5122 |
|  | (09183)- Protein families: signaling and cellular processes | 1126 |

**Supplementary Table S3:** List of genes which are related to diosgenin in *T. terrestris*

| S. No. | Gene ID | Enzyme name | Abbreviation | Enzyme commission number | Function of genes | Regulation |
| --- | --- | --- | --- | --- | --- | --- |
| 1. | >transcript_11011_unigene_10870_cds_4484 | Hydroxymethylglutaryl-CoA reductase | *HMGR* | EC:1.1.1.34 | Responsible for development stages. | Up-regulated in the all parts. |
| 2. | >transcript_101423_unigene_82578_cds_57949 | Mevalonate kinase | *MVK* | EC:2.7.1.36 | Essential in cell growth, maturation, and formation. | No expression |
| 3. | >transcript_65481_unigene_59157_cds_35079 | Mevalonate diphosphate decarboxylase | *MVD* | EC:4.1.1.33 | Important role in the regulating growth, sporulation, and stress tolerance. | Up-regulated in the all parts but down-regulated in leaf. |
| 4. | >transcript_28120_unigene_27514_cds_12458 | 1-deoxy-D-xylulose-5-phosphate synthase | *DXD* | EC:2.2.1.7 | Involved in regulation of biosynthesis and increased in steroidal hormones. | Up-regulated in the all parts. |
| 5. | >transcript_115207_unigene_90670_cds_66486 | 1-deoxy-D-xylulose-5-phosphate reductoisomerase | *DXR* | EC:1.1.1.267 | Play key role in localization into chloroplast and plastid of leaf cells. | Up-regulated in the all parts. |
| 6. | >transcript_33806_unigene_32889_cds_15450 | Isopentenyl-diphosphate delta-isomerase | *IDI* | EC:5.3.3.2 | Involved in cytosol/ER, peroxisomes, mitochondria, and plastid. | Up-regulated in the all parts. |
| 7. | >transcript_78673_unigene_68239_cds_43581 | Geanylgeranyl diphosphate synthase | *GPPS* | EC:2.5.1.29 | Main role in flower pollinators. | Up-regulated in the all parts. |
| 8. | >transcript_64263_unigene_58318_cds_34332 | Squalene epoxidase | *SQE* | EC:1.14.14.17 | Involved in developmental stages. | Up-regulated in the all parts. |
| 9. | >transcript_65063_unigene_58875_cds_34853 | 7-dehydrocholesterol reductase | *DWF7* | EC: 1.3.1.21 | Key role in brassinosteroid biosynthesis. | Up-regulated in the all parts. |
| 10. | >transcript_47375_unigene_45048_cds_23475 | Sterol 24-C methyltransferase | *SMT1* | EC: 2.1.1.41 | Important role in root growth, sterility | Up-regulated in the all parts. |
| 11. | >transcript_54100_unigene_50584_cds_27729 | Delta 14-sterol reductase | *FK* | EC 1.3.1.70 | Important role in cell division, embryogenesis, and development. | Up-regulated in the all parts. |
| 12. | >transcript_103639_unigene_83903_cds_59474 | Squalene synthase | *SQS* | EC: 2.5.1.21 | Essential role in growth development, and improve infertility. | Up-regulated in the all parts. |
| 13. | >transcript_6007_unigene_5926_cds_2619 | Phosphomevalonate kinase | *PMK* | EC:2.7.4.2 | Important role in regulation of terpenoids metabolism. | Up-regulated in the all parts. |
| 14. | >transcript_31524_unigene_30737_cds_14246 | Hydroxymethylglutaryl-CoA synthase | *HMGS* | EC: 2.3.3.10 | Key role in phytosterols biosynthesis. | Up-regulated in the all parts. |
| 15. | >transcript_35714_unigene_34669_cds_16543 | Housekeeping gene | *Alpha-* tubulin | - | Significant role in germination of pollen and tube growth, transport of vesicles and cell division. | Up-regulated in the all parts. |

**Supplementary Table S4:** Primer Identification of internal control genes and some of these genes are used for validation of qRT-PCR analysis of *T. terrestris*

| **Target genes** | **Transcript_Id** | **Forward/ Reverse** | **Primer sequences (5′ to 3′)** |
| --- | --- | --- | --- |
| Acetyl-CoA acetyltransferase  **HMGR** | transcript_11011_unigene_10870_cds_4484 | F | **TGTCACTTCCGTCCAAATCC (Sense)** |
|  |  | R | **AAGGAGGTGAATGCGGTATG (Antisense)** |
| Hydroxymethylglutaryl-CoA synthase  **MVK** | transcript_101423_unigene_82578_cds_57949 | F | **GGAGGCTGTGTTCTGACATT (Sense)** |
|  |  | R | **CCACCAGTCCCAGCAATAAA (Antisense)** |
| Hydroxymethylglutaryl-CoA reductase  **MVD** | transcript_57421_unigene_53200_cds_29985 | F | **CTGGAGGTAGGTACCAGAATTG (Sense)** |
|  |  | R | **CACGTGAAACTTGTCCCAATC (Antisense)** |
| Mevalonate kinase  **IDI** | transcript_33806_unigene_32889_cds_15450 | F | **AAGGCCATGTTCCAGGATAAA (Sense)** |
|  |  | R | **CCGTTTCTCCCATGTACTTCTC (Antisense)** |
| Phosphomevalonate kinase  **DXS** | transcript_28120_unigene_27514_cds_12458 | F | **GCTCAGTTTCTTGCGCTTTC (Sense)** |
|  |  | R | **ACTCTAGCAGCAATGTGAGATG (Antisense)** |
| Isopentenyl pyrophosphate isomerase  **DXR** | transcript_115207_unigene_90670_cds_66485 | F | **GGCACAATGACTGGAGTTCTTA (Sense)** |
|  |  | R | **CCAGTTCCTCTTGGTGCTTATC (Antisense)** |
| Farnesyl pyrophosphate synthase  **GPPS** | transcript_103896_unigene_84044_cds_59611 | F | **CAGAAGGGATCGAGAATCAGAAG (Sense)** |
|  |  | R | **GTGGGTACTCTGCCACAATAG (Antisense)** |
| Squalene synthase  **SQE** | transcript_112445_unigene_89105_cds_64946 | F | **TGGTGTTGGCCGTTTACTT (Sense)** |
|  |  | R | **GCCTAACTCCTTCTGCTTTGA (Antisense)** |
| Squalene epoxidase  **DWF7** | transcript_185181_unigene_140593_cds_101815 | F | **GAGCGTGATTTCTGATCATCCA (Sense)** |
|  |  | R | **GTGTTCCCGCCTCACTAATAC (Antisense)** |
| **HMGS** | transcript_34694_unigene_33731_cds_15932 | F | **GATATCTACTCCCTCGCTCTCA (Sense)** |
|  |  | R | **AGACCGATTTGACGGACTTG (Antisense)** |
| **SQS** | transcript_29196_unigene_28533_cds_12992 | F | **ATAGCCGACTACGACCTCTAC (Sense)** |
|  |  | R | **TGGAGTAGGAGACCAAGAGAAT (Antisense)** |
| **SMT1** | transcript_47375_unigene_45048_cds_23475 | F | **CTACCTCGCACACAAGATGAA (Sense)** |
|  |  | R | **GTAACGTTGACACCAGCAAAC (Antisense)** |
| **CYP51** | transcript_33990_unigene_33070_cds_15566 | F | **CTTACCTCCAACTGCAGACTATC (Sense)** |
|  |  | R | **GTCGTGAGAACAGACTCGAATAA (Antisense)** |
| **FK** | transcript_54100_unigene_50584_cds_27730 | F | **GCAGGAGAACTGCAAAGTAGA (Sense)** |
|  |  | R | **GCCCAGGATGACTCCTATTTC (Antisense)** |
| *Alpha-tubulin* | transcript_35714_unigene_34669_cds_16543 | F | **GGTCACTACACTATCGGAAAGG (Sense)** |
|  |  | R | **TCTTCGTCTTCCACTCCTTTG (Antisense)** |

**Supplementary Table S5:** Transcriptome data analysis SSRs or microsatellite data of *T. terrestris*

| **SSR data** | **Number of *T. terresris*** |
| --- | --- |
| Total number of sequences examined | 245069 |
| Total size of examined sequences (nt) | 179930901 |
| Total number of identified SSRs | 17623 |
| Numbers of SSR containing sequences | 15551 |
| Numbers of sequences containing more than 1 SSR | 1739 |
| Number of SSRs present in compound formation | 722 |
| Di-nucleotide | 9292 |
| Tri-nucleotide | 6580 |
| Tetra-nucleotide | 146 |
| Penta-nucleotide | 4 |
| AC/GT | 770 |
| AG/CT | 7842 |
| AT/AT | 1492 |
| CG/CG | 21 |
| AAC/GTT | 380 |
| AAG/CTT | 3065 |
| AAT/ATT | 335 |
| ACC/GGT | 407 |
| ACG/CGT | 175 |
| ACT/AGT | 90 |
| AGC/CTG | 694 |
| AGG/CCT | 341 |
| ATC/ATG | 1203 |
| CCG/CGG | 445 |
| AAAC/GTTT | 24 |
| AAAG/CTTT | 105 |
| AAAT/ATTT | 85 |
| AACC/GGTT | 5 |
| AAGC/CTTG | 6 |
| AAGG/CCTT | 6 |
| AAGT/ACTT | 1 |
| AATC/ATTG | 6 |
| AATG/ATTC | 6 |
| AATT/AATT | 18 |
| ACAG/CTGT | 5 |
| ACAT/ATGT | 20 |
| ACCG/CGGT | 1 |
| ACGG/CCGT | 1 |
| ACTC/AGTG | 11 |
| ACTG/AGTC | 1 |
| AGAT/ATCT | 22 |
| AGCC/CTGG | 1 |
| AGCG/CGCT | 2 |
| AGGC/CCTG | 1 |
| ATCC/ATGG | 1 |
| ATGC/ATGC | 16 |
| AAAAT/ATTTT | 1 |
| AAACG/CGTTT | 1 |
| AAATC/ATTTG | 4 |
| AACGG/CCGTT | 1 |
| AATGT/ACATT | 1 |
| ACACC/GGTGT | 6 |
| ACAGC/CTGTG | 1 |
| ACCAG/CTGGT | 1 |
| ACGAT/ATCGT | 1 |
| ACTGC/AGTGC | 1 |
| AGCTC/AGCTG | 1 |
